# Genetic architecture of phenological, morphological, and phytochemical traits in *Cannabis* landraces

**DOI:** 10.1101/2025.08.11.669724

**Authors:** Mehdi Babaei, Davoud Torkamaneh

## Abstract

Despite its long history of cultivation and diverse applications, *Cannabis sativa* remains underexplored at the genomic level, particularly in landrace populations that harbor untapped genetic diversity. In this study, we investigated the genetic architecture of 145 Iranian cannabis landrace accessions, including both male and female plants, using 233K common SNPs and genome-wide association studies (GWAS). Our analysis revealed three genetically distinct subpopulations shaped by geography, climate, and traditional cultivation practices. We identified 91 significant genomic regions associated with 40 phenological, morphological, and phytochemical traits, including 15 key loci with pleiotropic effects linked to multiple traits, including flowering time, plant architecture, biomass accumulation, and cannabinoid biosynthesis. These findings highlight the complex interplay between developmental and metabolic pathways in cannabis. The high heritability of most traits and rapid linkage disequilibrium decay underscore the potential of these landraces for high-resolution mapping and genetic improvement. This work provides a valuable genomic resource for marker-assisted selection, supporting the development of improved cultivars with tailored cannabinoid profiles and agronomic traits.

## 1. Introduction

Cannabis (*Cannabis sativa* L.), an annual flowering plant belonging to the Cannabaceae family, stands as one of the earliest domesticated crops in human history, with its origins tracing back approximately 3000-8000 years to East/Central Asia (Ahmed et al. 2008; Srinivasababu 2014; Small 2017; Crocq 2020; Schilling et al. 2020; Ren et al. 2021; Babaei et al. 2022, 2025). Throughout millennia, cannabis has been extensively cultivated and utilized for diverse applications, including fiber production, oil extraction, and its distinct medicinal and psychoactive properties (Anwar et al. 2006; Russo et al. 2008; Warf 2014; Barcaccia et al. 2020; Sun 2023). Additionally, early Iranian medical texts authored by physicians, such as Rhazes–the discoverer of alcohol and sulfuric acid (854–925) and Avicenna (980–1037), further attest to the therapeutic use of cannabis (Mahdizadeh et al. 2015; Bachir et al. 2022). This rich history has led to a wide array of genetic and phenotypic diversity across its indigenous populations, or landraces, which have adapted to various environments and human selection pressures (Soorni et al. 2017; Babaei and Ajdanian 2020; Babaei et al. 2024, 2025).

Despite its historical significance and versatile applications, scientific research into cannabis genetics and trait inheritance has lagged compared to other major crops, largely due to decades of prohibition and clandestine breeding practices (Mudge et al. 2018; Peng and Shahidi 2021; Torkamaneh and Jones 2021; Halpin-McCormick et al. 2024). This has also created a genetic bottleneck, leading to a reduction in diversity, particularly skewed towards drug-type varieties (Naim-Feil et al. 2021, 2022, 2023; Lapierre et al. 2023a; de Ronne et al. 2024; de Ronne and Torkamaneh 2025). However, with shifting legislation and increasing legalization globally, the cannabis market is experiencing unprecedented growth, driving a pressing need for a deeper understanding of its fundamental biology and genetic architecture (Hammond 2021; Statista 2024).

Cannabis is predominantly dioecious (male XY, female XX), with a diploid genome (2n=20), although monoecious forms and hermaphroditism can occur (Carpentier et al. 2012; Razumova et al. 2016; Braich et al. 2020; Monthony et al. 2024). Its taxonomy remains a subject of ongoing debate, with perspectives ranging from a single species (*C. sativa*) to multiple distinct species (*C. sativa*, *C. indica*, and *C. ruderalis*) (Small and Beckstead 1973; Anwar et al. 2006; Flores-Sanchez and Verpoorte 2008; McPartland 2018; Lapierre et al. 2023b). Modern classifications often extend beyond these taxonomic definitions, incorporating legal status (hemp vs. drug-type based on Δ^9^-tetrahydrocannabinol (THC) concentration < 0.3% and THC:CBD ratio > 1, respectively), phytochemical profiles (chemotypes; CBD, cannabigerol (CBG), cannabichromene (CBC), and cannabinol (CBN)), ecological adaptation (ecotypes), and biological characteristics (biotypes) (De Meijer and Hammond 2005; Sawler et al. 2015; Small 2015; Cherney and Small 2016; Babaei and Ajdanian 2020; Jang et al. 2020; McPartland and Small 2020; Lapierre et al. 2023b).

Recent policy shifts have expanded access to genomic and transcriptomic data and streamlined regulations for cannabis research (Hesami et al. 2020, 2024; Grassa et al. 2021; Monthony et al. 2024). This has catalyzed the adoption of advanced sequencing technologies, such as next-generation sequencing (NGS), which offer cost-effective, high-throughput data generation. Coupled with sophisticated bioinformatic tools, these innovations have greatly expanded the scope of genotype–phenotype association studies across diverse crop species (Torkamaneh et al. 2016, 2018; De Ronne et al. 2023; Satam et al. 2023; Abdi et al. 2024; Ahmed 2024). A critical challenge in cannabis breeding lies in accurately evaluating the genetic contributions from diverse or exotic and indigenous germplasms. This difficulty is compounded by the quantitative nature of many desirable traits and the significant environmental influence on plant performance, factors that have historically limited the widespread incorporation of such valuable genetic resources (Barcaccia et al. 2020; Ingvardsen and Brinch-Pedersen 2023). Over the past decade, genome-wide association studies (GWAS) have emerged as a powerful approach proving highly effective in dissecting the genetic basis of variation for intricate phenotypes, including plant physiological and agronomic characteristics (Belzile and Torkamaneh 2022). GWAS is now recognized as a premier tool for pinpointing genetic markers linked to traits of interest, particularly excelling where conventional methods fall short for complex traits (Alqudah et al. 2020; Belzile and Torkamaneh 2022; Torkamaneh and Belzile 2022).

In cannabis, flowering time is a critical agronomic trait, influenced by photoperiod (short-day plant) and strong genetic control (Lisson et al. 2000; Amaducci et al. 2008; Zhang et al. 2021; Babaei et al. 2022, 2024). Flowering directly impacts biomass accumulation, fiber quality, and cannabinoid production, with cannabinoids rapidly accumulating during early flowering stages (Amaducci et al. 2005, 2008; Salentijn et al. 2015; Petit et al. 2020a, b). Recent genetic studies have advanced our understanding of cannabis flowering, identifying key loci like *Autoflower1* and *Early1* (Toth et al. 2022), and *Autoflower2* (*FT1*) (Dowling et al. 2024) alongside multiple QTL associated with photoperiod sensing, circadian rhythms, and hormone signaling (Petit et al. 2020a). Beyond flowering, GWAS has been instrumental in elucidating genetic markers linked to a wide spectrum of cannabis traits, encompassing sex determination, cannabinoid biosynthesis pathways, fiber characteristics, and key morphological and agronomic features, and stress resilience (Petit et al. 2020c, a; Welling et al. 2020; Toth et al. 2022; de Ronne et al. 2023; Steel et al. 2023; Sun et al. 2023).

However, a key limitation in cannabis GWAS to date, particularly those using commercial cultivars, is their restricted genetic diversity. While valuable for specific traits, such studies often miss the broader genetic variation in landraces crucial for comprehensively dissecting complex traits. For instance, recent studies by de Ronne et al. (2024) on morphological traits and de Ronne and Torkamaneh. (2025) on cannabinoid profiles, both conducted on Canadian commercial cultivars, provided comprehensive catalogs of genetic variants but were inherently limited by the narrower diversity of cultivated lines compared to landraces.

This study aims to address these gaps by providing a comprehensive characterization of 145 Iranian cannabis landrace accessions. We employed high-density genotyping-by-sequencing (HD-GBS) to generate a catalog of 233,624 high-quality SNPs and GWAS to identify significant genetic loci and putative candidate genes associated with 42 key phenological, morphological, and phytochemical traits. Our findings will contribute significantly to the fundamental understanding of cannabis biology and provide valuable genetic resources and tools for accelerating molecular breeding efforts.

## 2. Materials and methods

### 2.1 Plant materials

Cannabis seed samples from 25 native populations (Table S1) of Iran were sourced from local markets and native farmers in various regions, ensuring genetic diversity and regional adaptation. The regions from which the seeds were collected encompassed five defined climatic zones based on the Köppen-Geiger climate classification method, with geographic latitudes between 25° and 40° North and Longitudes 45° and 65° East. Prior to cultivation, preliminary germination tests were conducted to evaluate key parameters.

### 2.2 Experimental design and growing conditions

Twenty seeds from each collected population were planted (Babaei et al. 2024). Thirty days after sowing (DAS), the seedlings were transferred to growth bags with a mixture consisting of garden soil, leaf mold, sand, and perlite in a 3:1:1:1 proportion. Based on the study by Amaducci et al. (2008), the planting was done 60 days before the photoperiod switch-off in Mashhad, Khorasan, Iran when the days start to shorten (the day length at sowing was 13 hours and 17 minutes) to ensure that the plants would have an adequate vegetative period before entering the reproductive phase. The plants were arranged in a randomized complete block design (RCBD) with three blocks and five observations, totaling 375 plants (50% male and 50% female), in the greenhouse complex of Ferdowsi University of Mashhad, Iran (36^°^16ʹN and 59^°^36ʹE with an altitude of 985 m). According to Spatial Analysis (SA), each unit, measuring 60 square meters, was divided into 25 rows and 15 columns, where each block included 5 columns. The aisles within each unit were arranged in order to place 16 plants per square meter. At the beginning of the reproductive stage (appearance of solitary flowers), male and female plants were separated and transferred to a unit with similar conditions, reducing the number of plants per square meter by half. The growing conditions and all cultural practices were previously described in Babaei et al. (2024).

### 2.3 Phenotypic data collection

Phenological and morphological data for this study were primarily derived from a previously published comprehensive phenotypic characterization of these Iranian cannabis landraces (Babaei et al. 2024). In that prior work, 12 phenological traits (covering both vegetative and reproductive stages) and 14 morphological traits were recorded on an individual basis for all 375 plants. For the current GWAS analysis, we specifically utilized six reproductive phenological stages and 12 morphological traits from this extensive dataset. A complete list of all phenological, morphological, and phytochemical (cannabinoid) traits, along with their defined categories, is provided in Table S2.

#### 2.3.1 Phenological traits

Based on the detailed phenological descriptor developed in our previous study (Babaei et al. 2024), phenological traits were recorded and defined for both male and female plants. For the present study, the focus was specifically on six reproductive stages. These traits, measured in days after sowing (DAS), include: 1) GV Point (GVP): Transition of bud phyllotaxis from opposite to alternate on main stem, marking entry into the reproductive phase (minimum 0.5 cm distance between alternate leaf petioles); 2) Start Flower Formation Time in Individuals (SFFI): First bud (solitary flower) appearance on individual plants (bell-shaped or closed sepals for male; symmetrical calyx with two styles for female); 3) Start Flower Formation Time in 50% Population (SFFP): First flower appearance in 50% of the population; 4) Start 10% Flowering Time in Individuals (SF10I): Minimum 10% of main inflorescence formed on individual plants; 5) Start 10% Flowering Time in 50% Population (SF10P): 10% of inflorescence formed in 50% of the population; 6) Flowering Time 50% in Individuals (FT50I): 50% of the main inflorescence formed on individual plants.

#### 2.3.2 Morphological traits

A total of 21 morphological traits were assessed in this study, categorized as Node and Branching Architecture, Growth and Structural Dimension, and Biomass Yield. These measurements were conducted at the end of the cultivation period, coinciding with the harvest of both male and female plants (Babaei et al. 2024). A schematic illustration of these key morphological traits is provided in Fig. 1a. Node and Branching Architecture traits included: 1) Number of Nodes to the Main Inflorescence (NTMI); 2) Number of Nodes to the First Lateral Shoot (NTFIS); 3) Number of Nodes on the main stem on Harvest day (NNH); 4) Number of Nodes on the Main Inflorescence (NMI); 5) Number of Lateral Shoot (NLS).

**Fig. 1.**
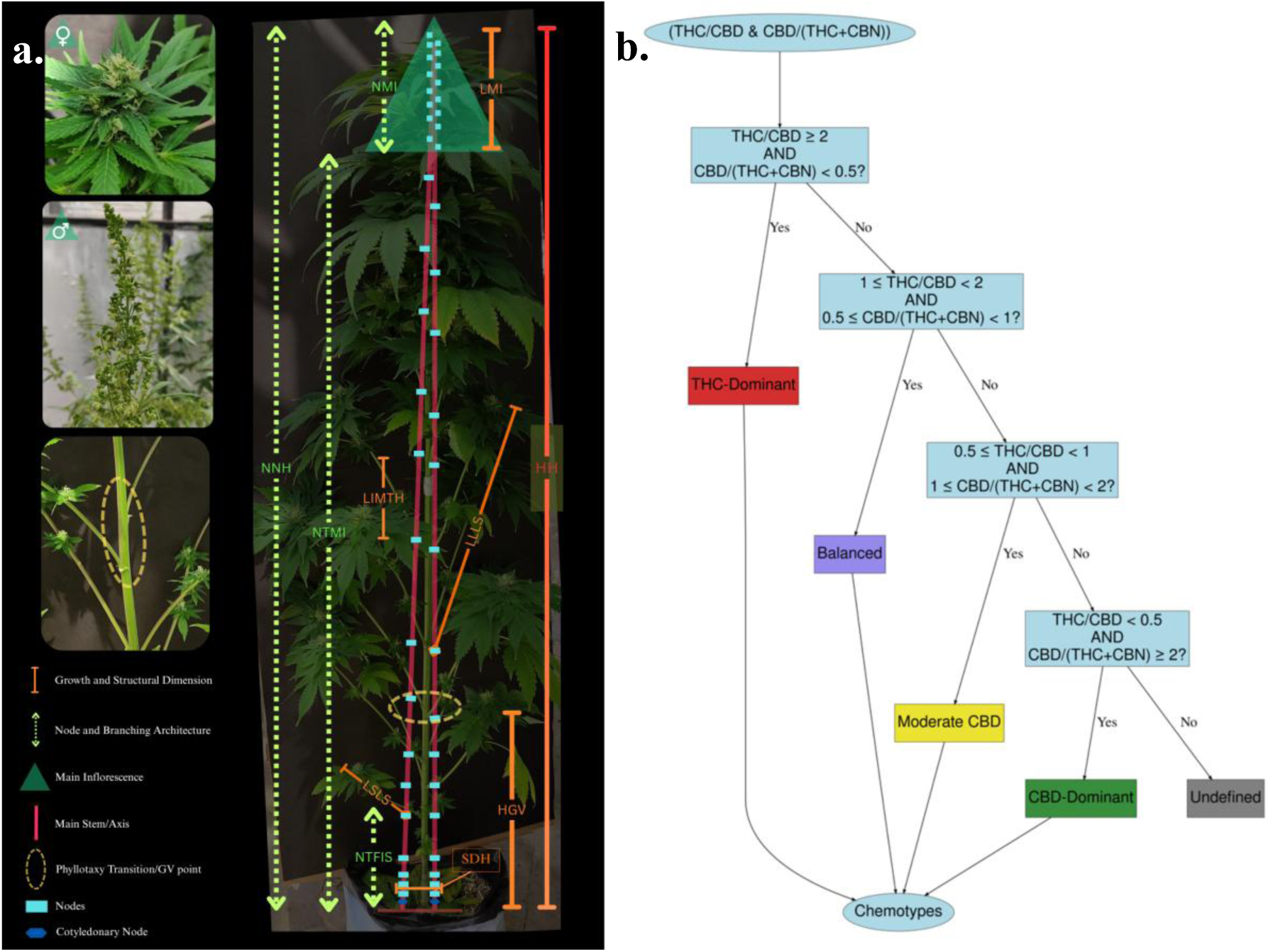
Visual representation of studied traits and chemotype classification in cannabis landrace accessions (**a**) Schematic illustration of key morphological traits measured in cannabis plants. This diagram highlights the various plant architectural and growth parameters assessed in this study (**b**) Decision tree for cannabis chemotype classification based on cannabinoid ratios. This flowchart illustrates the hierarchical classification of cannabis chemotypes using predefined thresholds for THC/CBD and CBD/(THC+CBN) ratios. Different pathways lead to the categorization of samples as THC-Dominant, Balanced, Moderate CBD, CBD-Dominant, or Undefined chemotypes. Abbreviations; NTMI: Number of Nodes to the Main Inflorescence, NTFIS: Number of Nodes to the First Lateral Shoot, NNH: Number of Nodes on the main stem on Harvest day, NMI: Number of Nodes on the Main Inflorescence, LSLS: Length of Shortest Lateral Shoot, LMI: Length of Main Inflorescence, LLLS: Length of Longest Lateral Shoot, LIMTH: Length of Internode in the Middle Third of the main stem on Harvest day, HH: Height on Harvest day, HGV: Height to GV Point, SDH: Stem Diameter on Harvest day.

The NNH was calculated as the sum of nodes on the main inflorescence and nodes to the main inflorescence (eq. 1). The NLS was determined based on the number of nodes to the main inflorescence and the first lateral shoot (eq. 2):

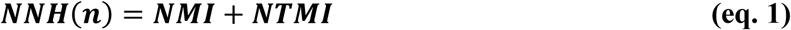

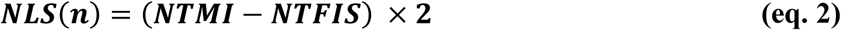

Growth and Structural Dimension traits included: 1) Length of Shortest Lateral Shoot (LSLS); 2) Length of Main Inflorescence (LMI); 3) Length of Longest Lateral Shoot (LLLS); 4) Length of Internode in the Middle Third of the main stem on Harvest day (LIMTH); 5) Height on Harvest day (HH); 6) Height to GV Point (HGV); 7) Relative Growth Rate (RGR); 8) Stem Diameter on Harvest day (SDH).

Traits such as LMI, HH, LIMTH, and HGV were measured using a tape measure with 0.01 m precision. SDH was measured at 5 cm above the soil surface using calipers with an accuracy of 0.01 mm. The RGR was also calculated using the following equation (eq. 3) and according to mg.g^-1^.day^-1^:

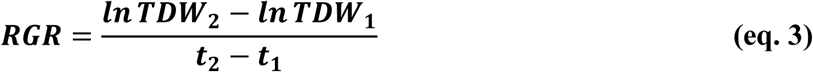

In this equation, the biomass increases from TDW_1_ to TDW_2_, on average during a time interval of t_1_-t_2_. *TDW_2_* and *TDW_1_* are the dry weights of plants at the time of harvest and in the seedling transplantation stage, respectively. Consequently, *t_2_* and *t_1_* represent the time of harvest and seedling transfer, respectively. *1n* signifies the Natural Logarithm (Babaei et al., 2024; Hoffmann and Poorter, 2002).

Biomass Yield traits included: 1) Dry Weight of Flowers (DWF); 2) Fresh Weight of Flowers (FWF); 3) Fresh weight of leaves (FWL); 4) Dry weight of leaves (DWL); 5) Fresh weight of stems (FWS); 6) Dry weight of stems (DWS); 7) Total Fresh Weight (TFW); 8) Total Dry Weight (TDW). A digital scale (AND-GF3000) with 0.001 gr accuracy was used for all fresh weight measurements of individual plant parts. For dry weight determination, samples were air-dried at room temperature (25°C) for 15 days before being re-weighed using the same digital scale. Total Fresh Weight (TFW) was calculated as the sum of fresh weights of stems, leaves, and flowers (eq. 4). Similarly, Total Fresh Weight (TFW) was calculated as the sum of fresh weights of stems, leaves, and flowers (eq. 5):

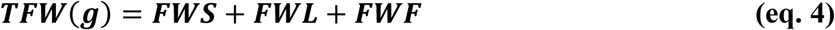

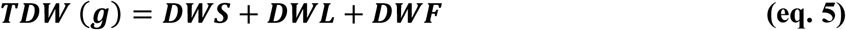

#### 2.3.3 Cannabinoid profiles

To prevent pollination, which is known to reduce cannabinoid content, male plants were separated from female plants prior to the onset of flowering (Lipson Feder et al. 2021). Sampling for cannabinoid analysis was conducted on female plants at the stage of flower maturity. This maturity was determined by daily visual inspection, specifically observing the browning of stigmas and the change in trichome color from clear to amber/brown, indicating peak cannabinoid production (Punja et al. 2023; Tran et al. 2025). Individual samples were collected from the main inflorescence of each plant as it reached this mature stage. Harvested samples were immediately placed in separate paper bags and air-dried in a dark environment at 25°C until completely dry. Subsequently, dried samples from each population within each experimental block were homogenized using a ceramic mortar.

##### Extraction procedure

First, 0.05 grams of dried plant tissue were extracted with 2 milliliters of methanol/chloroform (9:1 ratio). This mixture was then sonicated for 40 minutes in an ultrasonic bath, followed by centrifugation at 10,000 rpm for 15 minutes at 10°C. The supernatant was carefully separated and thoroughly dried by air (De Backer et al. 2009). The dried extracts were then solubilized in 2 mL of 80% methanol and sonicated for 10 minutes in an ultrasonic bath. After a final centrifugation for 5 minutes at 18,000g at room temperature, the supernatants were filtered through a 0.22 µm nylon filter and subsequently diluted by factors of 2 and 40 for analysis, following a modified method described by Mudge et al. (2018).

##### UPLC analysis

Biochemical analysis of these extracts was conducted at the Metabolomics Platform in the Institute of Nutrition and Functional Foods (INAF), Université Laval, Québec, QC, Canada. Cannabinoid analysis was performed using an Ultra Performance Liquid Chromatography (UPLC) Acquity H-Class system (Waters Corporation, Milford, MA, USA) coupled to an Acquity TUV detector (Waters Corporation, Milford, MA, USA). Compound separation was achieved on a Cortecs 1.6 µm, 2.1 mm X 150 mm (Waters Corporation, Milford, MA, USA) column, maintained at 30°C. The mobile phase consisted of (A) 20mM ammonium formate at pH 2.92 and (B) 100% acetonitrile. The gradient program was as follows: 0-6.4 minutes, 76% B; 6.5-8 minutes, 99% B. Conditions were reinitialized to 76% B from 8.1 to 10 minutes. The flow rate was 0.45 mL/min, and the injection volume was 2 µL. Detection was set at a wavelength of 228 nm. Cannabinoids were quantified using 5-point calibration curves prepared in the 1-100 mg/L range for all standards. The cannabinoid profiles quantified in this study, expressed as concentration (% w/w), included: Δ^9^-tetrahydrocannabinolic acid (THCA), Δ^9^-tetrahydrocannabinolic (Δ^9^-THC), cannabidiolic acid (CBDA), cannabidiol (CBD), cannabigerolic acid (CBGA), cannabigerol (CBG), cannabichromene (CBC), cannabinol (CBN), tetrahydrocannabivarin (THCV), cannabidivarin (CBDV), and Δ^8^-tetrahydrocannabinolic (Δ^8^-THC). It is important to note that Δ^8^-THC was consistently below the Limit of Detection (LOD) in all samples and was therefore excluded from subsequent analyses. To determine the Total THC Potential and Total CBD Potential, which represent the total decarboxylated forms, were calculated using the following equations (eq. 6 and eq. 7, respectively):

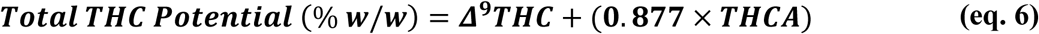

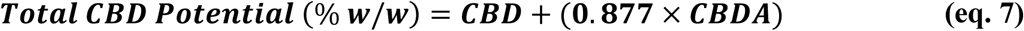

The coefficient 0.877, used in both equations, is derived from the molar mass ratio between the decarboxylated cannabinoids (Δ⁹-THC and CBD; 314.5 g/mol) and their acidic precursors (THCA and CBDA; 358.5 g/mol), accounting for the loss of the carboxyl group (CO₂) during decarboxylation.

Additionally, THC:CBD ratio and CBD:(THC+CBN) ratio were calculated to provide insights into the chemotype classification. For the first time, a novel chemotype classification system was developed in this study, integrating both the THC:CBD and CBD:(THC+CBN) ratios to define four distinct chemotype classes: THC-Dominant, Balanced, Moderate CBD, and High CBD. A decision tree illustrating this hierarchical classification based on predefined thresholds for these cannabinoid ratios is presented in Fig. 1b.

### 2.4 Genotyping and SNP calling

#### 2.4.1 Genotyping

Sampling was performed from healthy and young leaves of all individuals. Samples were then dried using the Freeze Dry System Alpha 2-4 LD plus for 24 hours. A total of 145 pooled samples, with each sample representing a unique combination of males and females from each population, were ground with metallic beads in a RETSCH MM 400 mixer mill (Fisher Scientific, MA, USA). DNA extraction was carried out using the Qiagen DNeasy® Plant Mini Kit. The extracted DNA quantity was assessed by a Qubit fluorometer with the dsDNA HS assay kit (Thermo Fisher Scientific, MA, USA), and their quality was randomly evaluated using agarose gel electrophoresis. DNA concentrations were adjusted to 10 ng/μl for all samples. Final DNA samples were used to prepare HD-GBS libraries with *BfaI* as described in Torkamaneh et al., (2021) at the Institut de biologie intégrative et des systèmes (IBIS), Université Laval, QC, Canada. Sequencing was conducted on an Illumina NovaSeq 6000 with 150 paired end reads at the Genome Quebec Service and Expertise Center (CESGQ), Montreal, QC, Canada.

#### 2.4.2 SNP calling and filtration

Sequencing data were processed with the Fast-GBS v2.0 using the *C. sativa* cs10 v2 reference genome (GenBank acc. no. GCA_900626175.2) (Grassa et al. 2018; Torkamaneh et al. 2020b). For variant calling a prerequisite of a minimum of 6 reads to call a single nucleotide polymorphism (SNP) was opted. After mapping against the cs10 v2 reference genome, an initial dataset of over 4.5 million variants was obtained. Raw SNP data were then filtered with VCFtools to remove low-quality SNPs (QUAL <10 and MQ <30) and variants with proportion of missing data exceeding 80% (Danecek et al. 2011). Following the first filtering step, which included quality filtering for minor allele frequency (MAF > 1%) and missing data > 80%, about 763 K variants were retained. A subsequent filtering step applying a minimum and maximum allele count of two reduced the dataset to ∼584 K variants. A second round of filtration was then applied, retaining only biallelic variants with heterozygosity less than 50% and a MAF of > 5%. Additionally, variants residing on unassembled scaffolds were removed. After these stringent filtering steps, the dataset was reduced to 233,624 high-quality SNPs on the scaffold, which were retained for downstream analysis (Table S3).

### 2.5 Genetic diversity analysis

#### 2.5.1 SNP-based genomic summary

Genetic diversity parameters were calculated across the entire panel (by chromosome, gender, and cluster/subpopulation). Read count and coverage were quantified from aligned reads using SAMtools (Danecek et al. 2021). MAF and heterozygosity level were determined using TASSEL v5.2.94 (Bradbury et al. 2007). Nucleotide diversity (*θπ*) was measured in 1kb sliding windows across the genome using the—*window-pi* option of VCFtools (Danecek et al. 2011). The proportion of SNPs located on annotated genes was determined by intersecting the filtered VCF file with a gene annotation BED file using BEDTools v2.31.1 (Quinlan and Hall 2010). Genome-wide gaps larger than 1 Mb were identified by subtracting the genomic regions covered by SNPs from the total chromosome lengths using BEDTools v2.31.1 (Quinlan and Hall 2010). To visualize the distribution of SNP density, a plot was produced with rMVP using the *plot.type = "d"* parameter, in combination with the gene density distribution (Yin et al. 2021). Additionally, variant types (e.g., multi-nucleotide polymorphisms (MNPs), insertions, deletions) and their predicted functional impacts (e.g., intergenic, intronic, synonymous, missense), along with the overall transition-to-transversion (Ts/Tv) ratio, were assessed using SnpEff v5.2e based on the *Cannabis sativa* reference genome (cs10 v2; GenBank accession: *GCA_900626175.2*) (Cingolani et al. 2012; Grassa et al. 2018).

#### 2.5.2 Linkage Disequilibrium (LD) decay and Haplotype Blocks (HBs)

Linkage Disequilibrium (LD) decay was assessed by calculating the squared allele frequency correlation (r^2^) using PopLDdecay v3.41 (Zhang et al. 2019). LD decay was calculated for the entire panel, for each chromosome, for each gender and for each genetic subpopulation, with a maximum physical distance of 500 kb (*-MaxDist 500*). Haplotype blocks (HBs) were identified using PLINK v1.90b5.3 by computing pairwise LD (r^2^), considering a window of 999kb SNPs ‘*– ld-window-kb 999’* (Purcell et al. 2007).

### 2.6 Population structure analysis

#### 2.6.1 Population structure inference

Population structure and admixture were determined using the variational Bayesian inference algorithm implemented in fastStructure v1.0 for K (number of subpopulations) values from 1 to 10 (Raj et al. 2014). The optimal K was estimated using the ChooseK tool, and admixture proportions visualized with Distruct v2.3. Discriminant Analysis of Principal Components (DAPC) was performed using the R package ‘*adegenet’* version 2.1.10 (Jombart et al. 2010). The optimal number of clusters (K) was estimated using the ‘*find.clusters*’ function, exploring up to 40 clusters, and determined by the minimal Bayesian Information Criterion (BIC). To visualize the DAPC using the ‘scatter’ function, the optimal number of principal components (PCs) (22 PCs) was estimated with two cross-validation procedures using ‘*optim.a.score*’ and ‘*xvalDapc’* (exploring PCs up to a maximum of 144) (Fig. S4a).

#### 2.6.2 Phylogenetic and kinship analysis

Phylogenetic relationships among accessions were inferred using the Neighbor-Joining (NJ) method in MEGA version 11 (Jin and Nei 1990; Kumar et al. 2018). This was based on genetic distances calculated from SNP data, using three different nucleotide substitution models: Jukes-Cantor (TC), Tajima-Nei (TN), and Maximum Composite Likelihood (MCL). A bootstrap consensus tree was formed from 1000 replicates. The resulting phylogenetic tree was edited and beautified using iTOL version 7.2 (Letunic and Bork 2021). Genetic distances were also estimated in MEGA version 11 using the p-distance model with Gamma Distributed (G) rates among sites (Gamma Parameter = 1) (Kumar et al. 2018). Pairwise distances, within-group average distances, between-group average distances, and overall mean distances were calculated. For all distance calculations, a Partial deletion approach was used for gaps/missing data with a Site Coverage Cutoff of 95%. Variance estimation for these distances was performed using 1000 bootstrap replicates. The pairwise genetic distances were subsequently visualized as a heatmap in R using the ‘*pheatmap’* package (Kolde and Kolde 2015; Team 2020). Kinship analysis was performed using TASSEL v5.2.94 with the Centered_IBS method, and the kinship matrix was plotted with GAPIT v3 (Bradbury et al. 2007; Wang and Zhang 2021).

### 2.7 Genome-wide association analysis

Genetic associations between markers and traits were investigated using the Bayesian-information and linkage-disequilibrium iteratively nested keyway (BLINK) method implemented in GAPIT v3 (Huang et al. 2019; Wang et al. 2022). This analysis focused on 233,624 high-quality SNPs and phenotypic data collected for 42 distinct traits. To mitigate the occurrence of false positive associations, the analysis incorporated both population structure (represented by a P matrix derived from fastStructure with K=3) and kinship (a K* matrix generated using TASSEL v5.2.94) as statistical covariates (Bradbury et al. 2007; Raj et al. 2014). A stringent significance threshold for marker-trait associations was established at a False Discovery Rate (FDR) < 0.05, which was achieved through adjustment using the Benjamini–Hochberg correction (de Ronne et al. 2024). Furthermore, markers explaining less than 3% of the Proportion of Phenotypic Variance Explained (PVE) were excluded from the final analysis, as they were deemed to provide limited informative value. Visual representations of the GWAS results included Manhattan plots, which depicted the –log_10_(*p*) distribution of markers across chromosomes (generated using rMVP with *plot.type* = "*m*"), and quantile–quantile (QQ) plots, created with GAPIT v3, to assess model fit (Wang and Zhang 2021; Yin et al. 2021). Additionally, boxplots illustrating the phenotypic effects of different allelic classes for significant markers were generated using the ‘*ggplot2’* package in R (Wickham et al. 2016; Team 2020). A comprehensive circular genomic map illustrating the genome-wide distribution of significant SNP markers and their associated traits was generated using shinyCircos-V2.0 (Wang et al. 2023).

### 2.8 Candidate gene identification and annotation

For the identification of putative candidate genes, haplotype blocks (HBs) were defined based on significant SNP markers identified through GWAS. Given the genetic diversity in cannabis, only markers exhibiting high linkage disequilibrium (r^2^ ≥0.75) with the significant SNPs were retained to delineate these HBs (de Ronne et al. 2024). Genes situated within these defined HBs, delimited by the 5’-most and 3’-most SNP positions of each block, were considered as putative candidate genes. Haploview v4.1 was used for visual inspection of marker positions within HBs, with the required input formats generated using PLINK’s *‘–recode HV*’ function (Barrett et al. 2005). Functional annotation and interpretation of these candidate regions were performed using SnpEff v5.2e, leveraging the *Cannabis sativa* reference genome (cs10 v2; GenBank accession: *GCA_900626175.2*) (Cingolani et al. 2012; Grassa et al. 2018). Additionally, Gene Ontology (GO) terms were extracted from the NCBI *Cannabis sativa* Annotation Release 100 to characterize the potential biological roles and pathways associated with these candidate genes.

### 2.9 Statistical analysis

The data were first evaluated using the Outlier Grubbs approach to detect and remove any outliers (Adikaram et al. 2015). Basic descriptive statistics for all traits were calculated to summarize their distribution and variability. One-way ANOVA was performed to assess the significance of trait variation among accessions and subpopulations, followed by post-hoc comparisons between clades at the 95% confidence level according to Tukey’s HSD test. Skewness and kurtosis analyses, assessing data normality and residual errors based on the distribution type, were conducted using Minitab^®^ Statistical Software 21 (Cain et al. 2017; Minitab 2021). Missing data was imputed using the Classification And Regression Tree (CART) method implemented in the ‘*AllInOne*’ package in R (Team 2020; Najafabadi et al. 2023). For phenological and morphological data, Spatial Analysis (AR1⊗AR1) was applied to account for potential spatial effects and minimize environmental impact, considering the experimental layout (15 columns and 25 rows). The data fitting approach considered population as a fixed factor and block as a random factor. The model’s performance was evaluated using four primary metrics: Root Mean Square Error (RMSE), Mean Squared Error (MSE), Normalized Root Mean Squared Error (NRMSE), and Coefficient of Variation (CV) (Babaei et al. 2024). Both broad-sense heritability (H^2^) and SNP-based heritability (h^2^) were estimated for all traits to quantify the genetic contribution to phenotypic variation. H^2^ was determined by the ‘*AllInOne*’ package and h^2^ was estimated using GCTA software with the restricted maximum likelihood (REML) method, based on the genomic relationship matrix (GRM) (Yang et al. 2011; Najafabadi et al. 2023). Pearson’s correlation coefficients were calculated to assess the interrelationships between all phenotypic traits, utilizing the package ‘*corrplot*’ packages in R (Wei et al. 2017). A heatmap of cannabinoid profiles was generated using the ‘*pheatmap’* package in R (Kolde and Kolde 2015). Principal Component Analysis (PCA) was conducted using the ‘*Factoextra’* package in R (Kassambara and Mundt 2017). Boxplots, violin plots, and the heatmap of the confusion matrix were generated using Python 3.12 with ‘*matplotlib’* and ‘*seaborn’* package (Bisong 2019).

## 3. Results

### 3.1 Phenotypic characterization of cannabis landraces

A comprehensive phenotypic analysis was conducted on 145 cannabis landrace accessions, comprising 72 females and 73 males, encompassing a wide range of 42 phenotypic and phytochemical traits. A comprehensive list of evaluated traits is provided in Table S2, with overall distributions (Fig. 2) and analysis of trait interrelationships through Pearson’s correlation coefficients (Fig. S1).

**Fig. 2.**
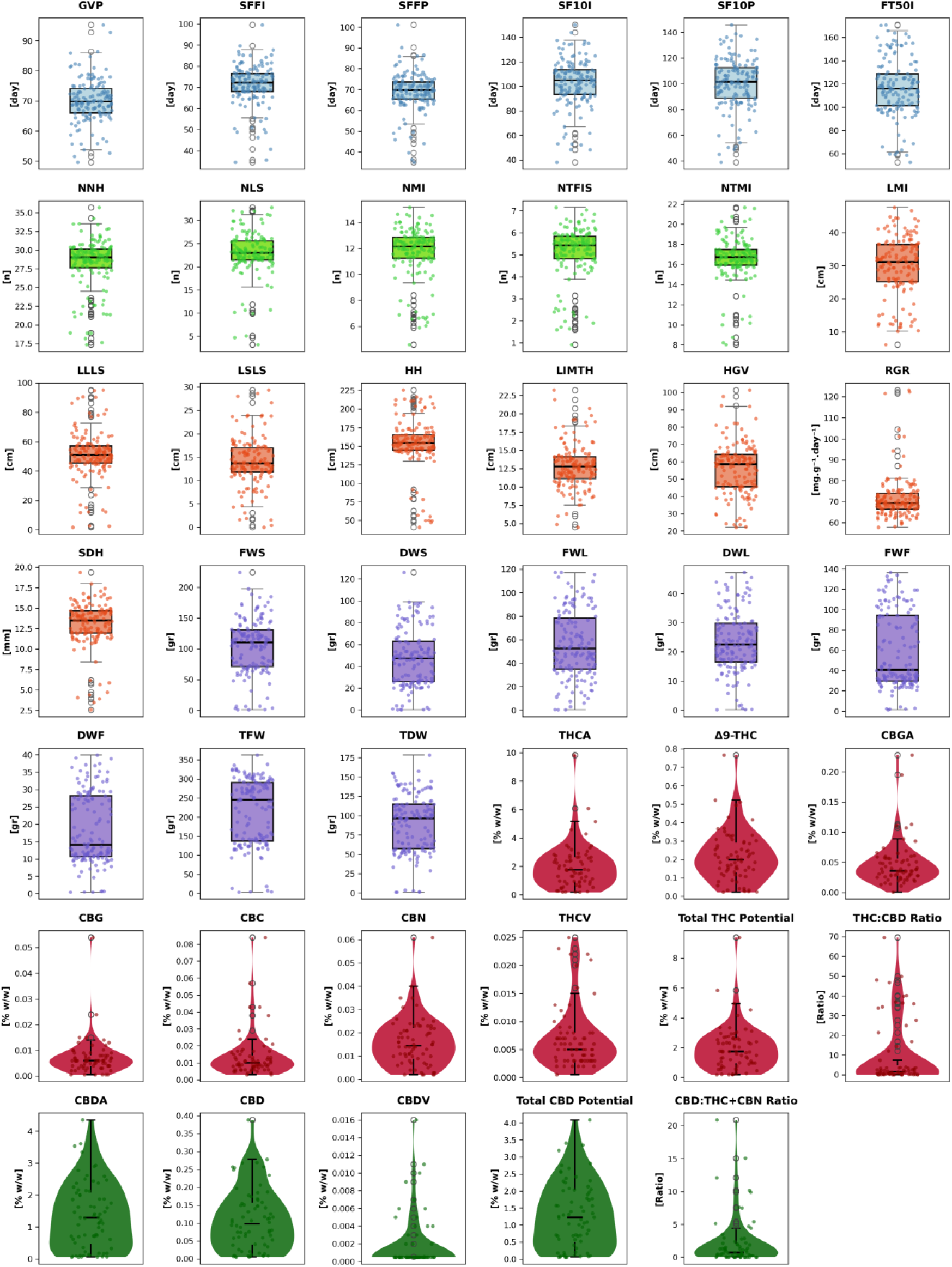
Distribution of phenological (blue), morphological (Node and Branching Architecture (light green), Growth and Structural Dimension (orange), and Biomass Yield (purple)), and cannabinoid profiles (THC-related (red) and CBD-related (dark green)) across 145 landrace accessions.

#### 3.1.1 Phenological traits

A comprehensive analysis of six phenological stages revealed significant variation in reproductive phase traits (Table S4 and Fig. 2). Traits exhibited significant differences, with GVP showing significance at P ≤ 0.01, and the remaining five stages (SFFI, SFFP, SF10I, SF10P, and FT50I) at P ≤ 0.001. Skewness and kurtosis analyses indicated non-normal distributions, particularly for SFFI and SFFP, suggesting tendencies towards earlier flowering and presence of extreme phenotypes. Furthermore, the H^2^ for these traits ranged from 0.41 (GVP) to 0.60 (FT50I).

#### 3.1.2 Morphological traits

The analysis of twenty-one morphological traits demonstrated considerable variation across the accessions (Table S5 and Fig. 2). All morphological traits exhibited significant differences among the accessions (P ≤ 0.001), except FWF and DWF, which were not statistically significant. High variability was observed in traits like LSLS (CV% 37.54) and TFW (range 359.95 g). The H^2^ for morphological traits varied widely, ranging from 0.41 (LIMTH) to 0.80 (TFW), suggesting a broad spectrum of genetic influence.

#### 3.1.3 Cannabinoid profiles

Analysis of fifteen cannabinoid profiles revealed substantial variability in accumulation among the landrace accessions (Table S6 and Fig. 2). Significant variation was observed for Δ^9^-THC (P ≤ 0.01), CBN (P ≤ 0.05), CBDV (P ≤ 0.01), Total THC Potential (P ≤ 0.05), and the CBD:(THC+CBN) ratio (P ≤ 0.001). Δ^8^-THC was undetectable in all samples. Individual cannabinoid concentrations and ratios showed high CV% values (e.g., CBG 99.98%, CBDV 164.5%), reflecting diverse chemotypes. Skewness and kurtosis analyses indicated non-normal distributions, often with high kurtosis, suggesting a deviation from normality characterized by a greater concentration of values in the tails. Heritability was generally high, with H^2^ ranging from 0.46 to 0.91 and h^2^ from 0.38 to 1.00. Total THC Potential (H^2^=0.90, h^2^=1.00) and Total CBD Potential (H^2^=0.74, h^2^=0.64) exhibited strong genetic control over primary cannabinoid biosynthesis.

### 3.2 Genotyping and population structure

#### 3.2.1 High-Density SNP coverage of the GWAS panel

Sequencing libraries generated ∼400 M reads, averaging approximately 2.7 M reads per sample. The final catalog of 233,624 (211,621 SNPs; 7,500 multi-nucleotide polymorphisms (MNPs), 6,588 insertions, and 7,915 deletions) common variants (MAF > 5%) displayed a density of 273.5 markers per Mb (one marker every ∼3.6 kb), with 20.8% of the genotypes being heterozygous and an average MAF of 12.6%. Notably, 21.2% of variants were located on annotated genes. Across the genome, fifteen gaps larger than 1 Mb were identified, with the largest being 2.3 Mb (Fig. S2a; Table S7). Additionally, a gender-based analysis revealed that males had a higher average MAF (13.61% vs. 11.48%), and heterozygosity (22.58% vs. 18.94%) compared to females (Table S8). Analysis of 233,624 variants revealed that 95.1% were modifiers with minimal functional impact, primarily in non-coding regions (intergenic (40.4%), intronic (11.4%), upstream (13.8%), and downstream (16.2%)). Among coding variants, 60.8% were synonymous and 38.7% missense, yielding a missense-to-silent ratio of 0.64. The overall transition-to-transversion (Ts/Tv) ratio was 1.76, indicating a balanced mutation spectrum and overall data quality.

LD decay across the analyzed chromosomes was compact, with the maximum r² values ranging from 0.42 (chr09) to 0.51 (chrX) (Fig. S2b and Table S7). On average, r^2^ dropped to half its maximum within 300 bp. The results from the LD and haplotype block (HB) analysis were congruent for male and female plants, with both sexes showing rapid LD decay and similar values for maximum r² (0.44 for females and 0.45 for males) (Fig. S2c and Table S8). However, slight differences were noted in the HB analysis. Male plants exhibited larger haplotype block spans (mean span of 0.51 kb and maximum span of 164.06 kb) compared to females (mean span of 0.43 kb and maximum span of 87.64 kb). Additionally, the proportion of SNPs in haplotype blocks was slightly higher in males (36.85%) compared to females (32.53%).

#### 3.2.2 Population structure, genetic diversity, and evolutionary dynamics

The analysis of the population structure using the full catalog of SNPs revealed three main subpopulations (K=3), supported by DAPC and model-based clustering by fastStructure (Fig. 3). Three main subpopulations were identified as K1 (clade I = 64 accessions), K2 (clade II = 64 accessions), and K3 (clade III = 17 accessions) (Table S9 and Fig. S3). DAPC and fastStructure analyses consistently supported K=3 as the optimal number of clusters, with high agreement in cluster assignments (Fig. 3a(ii), 3b; Table S9; Fig. S4). BIC comparisons revealed minimal differences between K=2 (1359.038) and K=3 (1359.258), with fastStructure identifying K=3 as the optimal population structure (Fig. S4b). Maximizing the marginal likelihood confirmed K=3 as optimal, revealing subpopulation diversity and admixture patterns (Table S9; Fig. 3a(iii); Fig. S5).

**Fig. 3.**
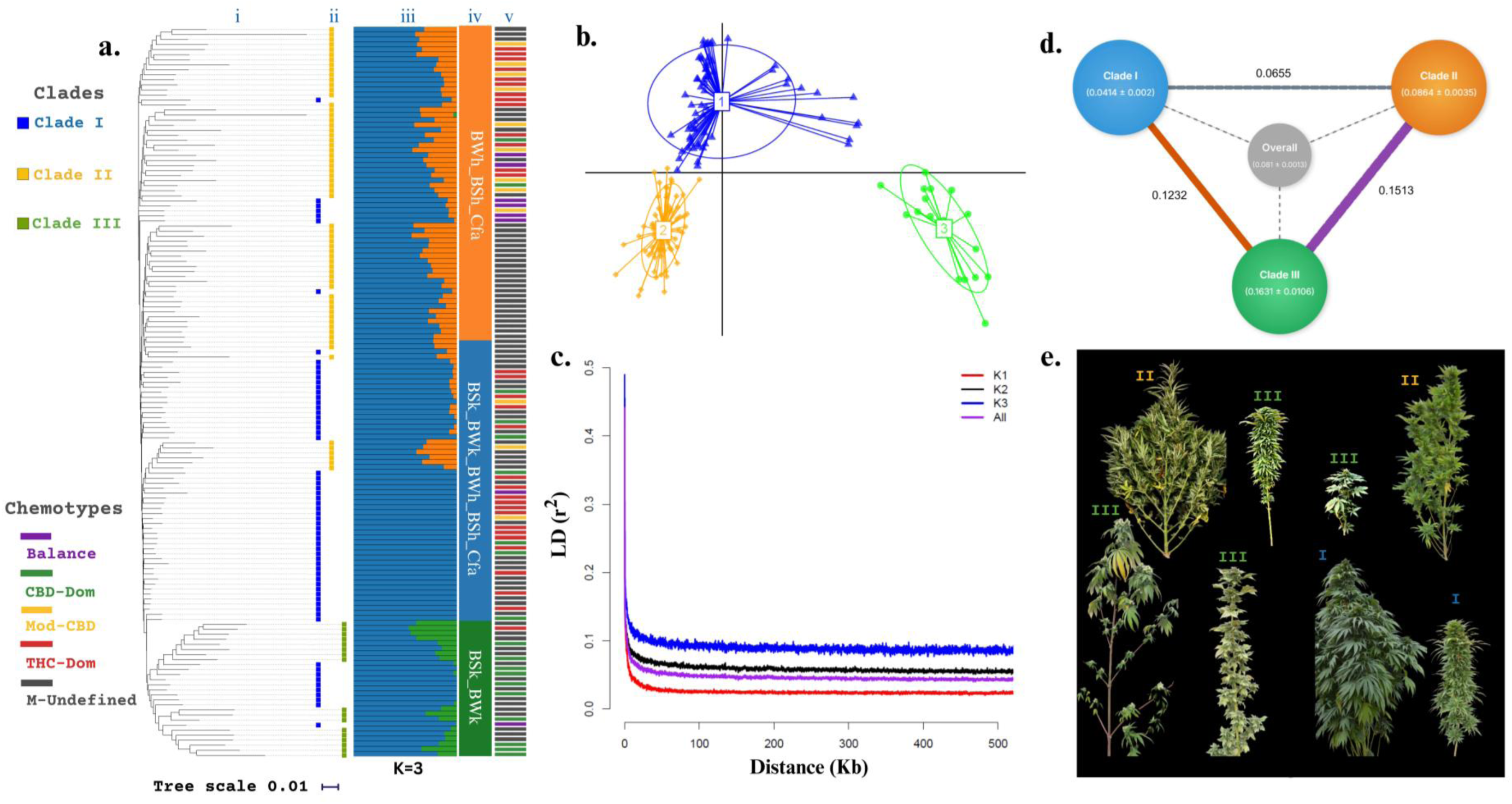
The population structure of 145 cannabis landrace accessions highlights three distinct genetic clusters (clade I, clade II, and clade III). (**a**) Comprises four sub-panels: (i) A phylogenetic tree highlighting the three clades. (ii) Discriminant analysis of principal components (DAPC), showing the clustering of accessions into the same clades. (iii) An admixture bar plot generated with fastStructure, representing proportional ancestry for each accession across the clades (K=3). (iv) Climate zones associated with each clade, corresponding to environmental regions where seeds were collected (BSh: arid, steppe, hot; BSk: arid, steppe, cold; BWh: arid, desert, hot; BWk: arid, desert, cold; Cfa: temperate, without dry season, sizzling summer), detailed in Fig. S6. (v) Chemotype classification of accessions based on cannabinoid ratios, detailed in Fig. 1b (**b**) The DAPC scatter plot demonstrates genetic variation and clustering of accessions into the three clades. (**c**) Linkage disequilibrium (LD) decay analysis shows LD patterns across the population and each clade. (**d**) Genetic distance diagram, visualizing within-group genetic distances for clades I, II, and III, and pairwise distances between them. A central node (gray) represents the overall genetic distance. (**e**) Representative plant images illustrating phenotypic diversity within and across the three clades.

Of the initial 233,624 variants, 192,465 sites were utilized in evolutionary and distance analyses, comprising 52,894 conserved sites and 139,571 variable sites. The phylogenetic tree (Fig. 3a(i)) aligned with DAPC (Fig. 3a(ii)) and fastStructure clustering (Fig. 3a(iii)), though with a different pattern: while clade I was identified as dominant subgroup in admixture analysis, some of its accessions were positioned between clades II and III in the phylogenetic tree (Fig. 3a(i)). Genetic distance analysis further supported the observed population structure. Within-group distances were lowest in clade I (0.0414 ± 0.002) and highest in clade III (0.1631 ± 0.0106), with clade II showing intermediate diversity (0.0864 ± 0.0035). Between-group distances were smallest between clades I and II (0.0655), and highest between clades II and III (0.1513), followed by clades I and III (0.1232) (Fig. 3d; Fig. S6).

Consistent with clustering results, genetic diversity metrics (Table S10 and Fig. 3c) and geographic distribution patterns (Fig. 3a(iv) and S7) revealed distinct characteristics among clades. Clade II, found in warmer regions (BWh, BSh, and Cfa climate zones), exhibited the highest genetic diversity ((*θ_π_*); 8.68×10^-4^) with MAF of 17.3% and heterozygosity of 29.8%, significantly exceeding values in clade I (*θ_π_*: 3.98×10^-4^; MAF=7%; heterozygosity=11.2%), which was present in across all climate zones. In contrast, clade III, mainly distributed in colder regions (BSk and BWk climate zones), showed intermediate genetic diversity (*θ_π_*: 7.66×10^-4^; MAF=14.5% and heterozygosity=22.6%). Clade II exhibited the highest HB density (34.4 HB/Mb) compared to clades I (15.4 HB/Mb) and III (6.8 HB/Mb). However, clade II’s HBs were shorter (mean span 0.34 kb) than those in clade I (1.38 kb) and clade III (0.81 kb) (Table S10). Despite this, the LD analysis revealed similar patterns across clades (Table S10 and Fig. 3c). Furthermore, kinship analysis revealed notable relationships within clade I, suggesting shared ancestral backgrounds within this subgroup (Fig. S8).

#### 3.2.3 Phenotypic insights from population structure

Phenotypic variation among the three genetic clades revealed distinct patterns across phenological, morphological, and phytochemical traits. Phenological analysis (Table S11) showed notable differences in reproductive phase initiation among clades (Fig. S9). Specifically, traits like GVP (P ≤ 0.01), SFFI (P ≤ 0.05), and SFFP (P ≤ 0.05) exhibited significant variation among clades. Clades I (e.g., SFFI: Mean = 70.87 ± 9.75 days, Range = 53.2 days) and clade III (e.g., SFFI: Mean = 65.95 ± 18.29 days, Range = 51.02 days) displayed wide flowering time ranges, encompassing both early and late accessions. In contrast, clade II (e.g., SFFI: Mean = 72.96 ± 4.88 days, Range = 25.21 days) consistently showed a narrower range (Table S11). This substantial phenological diversity was parallel by the observed morphological variations within and between clades (Table S12, Fig. S10 and Fig. 3e).

Notably, clades I and III displayed a higher phenotypic variation range for traits related to Node and Branching Architecture (e.g., NNH, NMI, NTFIS, NTMI all showing P ≤ 0.001) as well as Growth and Structural Dimension (e.g., LMI (P ≤ 0.001), LSLS (P ≤ 0.001), RGR (P ≤ 0.001), and SDH (P ≤ 0.01)) (Table S12). Specifically, clade III exhibited the lowest mean for NNH (23.04 ± 2.99 nodes), significantly fewer nodes than clades I (28.55 ± 3.56 nodes) and II (29.46 ± 1.54 nodes), aligning with the observation of longer internode lengths. For plant HH, clade III had the lowest mean (133.09 ± 58.84 cm) but the widest range (165.05 cm), indicating it indeed encompasses both the shortest and tallest plants (P ≤ 0.05). Interestingly, despite these significant structural and architectural differences, Biomass Yield traits (e.g., FWS, DWS, TFW, and TDW) generally showed no significant difference (Table S12). However, while FWF did not show significant differences, DWF exhibited significant differences (P ≤ 0.05) among clades (Fig. S10 and Table S12).

Most notably, the phytochemical profiles of these landraces aligned strongly with the established population structure and geographical origins (Table S13). Analysis of chemotype classes (THC-Dominant, Balanced, Moderate CBD and CBD-Dominant) distribution across clusters revealed a clear pattern: clade I exhibited a mixed chemotype profile, predominantly characterized by THC-dominant accessions, showing a mean Total THC Potential of 1.832 ± 1.079% and a lower mean Total CBD Potential of 1.172 ± 0.924%. Clade II comprised accessions showing either THC-dominant or moderate-CBD profiles, often displaying the highest mean concentrations for several cannabinoids, such as THCA (2.554 ± 2.01%) and Δ^9^-THC (0.269 ± 0.159%). Significantly, clade III was almost exclusively composed of CBD-dominant chemotypes (Table S13, Fig. 3a(v) and Fig. S11). This clade was characterized by significantly higher levels of CBD-related cannabinoids (e.g., CBDA, CBD, CBDV), with a markedly higher CBD:(THC+CBN) ratio (7.27 ± 7.124, P ≤ 0.001) compared to clades I and II. Conversely, Clade III showed significantly lower levels of THC-related cannabinoids (e.g., THCA, Δ^9^-THC, CBN, THCV, CBC), with Total THC Potential averaging 0.659 ± 0.652% (P ≤ 0.01) (Table S13). The heatmap and PCA of cannabinoid profiles consistently supported chemotype-based clustering, further reinforcing the observed distinctions among accessions (Fig. S12 and S13).

### 3.3 Genome-wide association study (GWAS) in cannabis landraces

A GWAS using BLINK model, which accounts for population structure and kinship, was conducted on 145 cannabis landrace accessions using 233,624 variants, identifying of 91 loci associated with 40 phenological, morphological, and phytochemical traits under a stringent threshold (*p*-value = 1.77×10^−7^, FDR <0.05). Since some loci were associated with multiple traits across or within trait categories, the counts per category reflect unique significant markers.

#### 3.3.1 Phenological Traits

GWAS for phenological traits revealed 10 significant markers associated with six distinct phenological stages, each exerting a measurable phenotypic effect, with PVE values ranging from 5.3% (Cs3 for GVP) to 44.7% (Cs8 for SFFP) (Fig. 4; Table 1; Fig. S14a). Key loci included a novel auto-flowering locus, designated *AutoFlower3* (*CsFT3*), which encompasses two closely linked markers: Cs1 (Chr8:7039513; PVE: 19.9% for GVP) and Cs5 (Chr8: 7038258, PVE: 37% for SSFI, 39.3% for SFFP, 25.6% for SF10I, 24.6% for SF10P and 20.9% for FT50I). Another important locus, designated *FloweringTime_Maturity4* (*CsFT4*), was identified on Chr7: 7258159 (Cs9) with PVE values of 13.3% for SF10I and 30.6% for FT50I. Additionally, *CircadianFloweringLocus1* (*CsCFL1*), located on Chr9:30838575 (CS2), showed strong associations with multiple traits (PVE: 33.2% for GVP, 41.4% for SF10I, 41.3% for SF10P and 24.3% for FT50I). The allelic impact of *CsCFL1* was particularly pronounced. For example, in the case of FT50I, individuals carrying the ‘CC’ allele exhibited a significantly higher mean flowering time (121 days) compared to those with the ‘AA’ allele (79 days), demonstrating clear allelic differentiation (Fig. S15). Furthermore, several significant markers were located on the X chromosome (e.g., Cs4, Cs8, Cs10), indicating its involvement in the regulation of phenological traits. Overall, these markers are of particular interest to cultivators aiming for faster crop turnover, as they are associated with shorter flowering or maturation times.

**Fig. 4.**
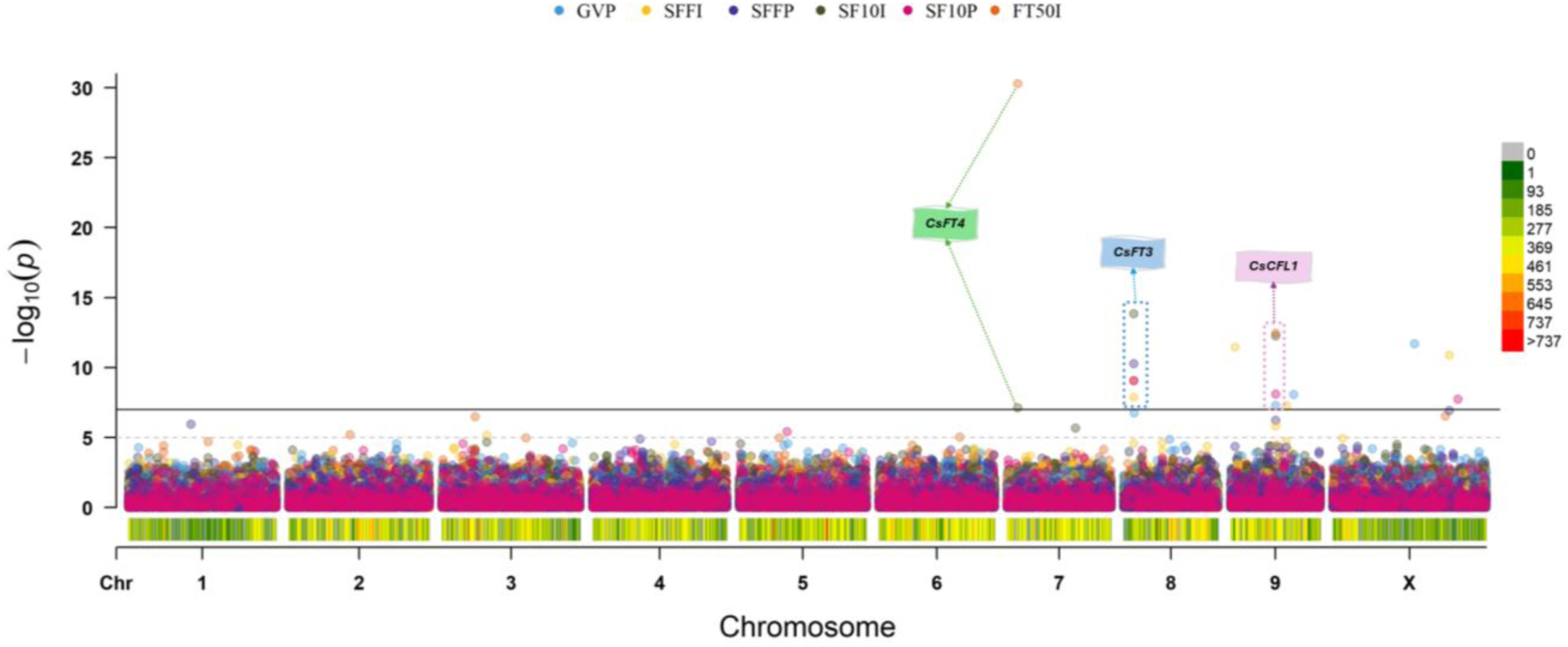
Manhattan plot illustrating GWAS results for phenological traits.

**Table 1.** List of 10 markers associated with six phenological stages.

| Traits | Marker ID | Locus ID<sup>a</sup> | Chr | Position (bp)<sup>b</sup> | *p*-value | MAF<sup>c</sup> (%) | Allele<sup>d</sup> | Effect<sup>e</sup> | PVE<sup>f</sup> (%) |
| --- | --- | --- | --- | --- | --- | --- | --- | --- | --- |
| GVP | Cs1 | *CsFT3* | 8 | 7039513 | 1.7E-07 | 9 | A/C | 5.04 | 19.9 |
|  | Cs2 | *CsCFL1* | 9 | 30838575 | 5.2E-08 | 5 | C/A | 7.56 | 33.2 |
|  | Cs3 | - | 9 | 43193302 | 8.7E-09 | 24 | G/A | 3.98 | 5.3 |
|  | Cs4 | - | X | 55838797 | 2.0E-12 | 6 | G/A | -8.30 | 24.6 |
| SFFI | Cs5 | *CsFT3* | 8 | 7038258 | 1.3E-08 | 8 | G/A | 6.75 | 37.0 |
|  | Cs6 | - | 9 | 3109357 | 3.5E-12 | 12 | C/T | -5.22 | 13.8 |
|  | Cs7 | - | 9 | 38458348 | 5.4E-08 | 7 | A/T | 5.21 | 5.6 |
|  | Cs8 | - | X | 79614615 | 1.3E-11 | 8 | C/T | 5.97 | 33.0 |
| SFFP | Cs5 | *CsFT3* | 8 | 7038258 | 5.2E-11 | 8 | G/A | 8.44 | 39.3 |
|  | Cs8 | - | X | 79614615 | 1.2E-07 | 8 | C/T | 5.53 | 44.7 |
| SF10I | Cs9 | *CsFT4* | 7 | 7258159 | 7.4E-08 | 26 | A/- | 9.95 | 13.3 |
|  | Cs5 | *CsFT3* | 8 | 7038258 | 1.4E-14 | 8 | G/A | 19.17 | 25.6 |
|  | Cs2 | *CsCFL1* | 9 | 30838575 | 5.7E-13 | 5 | C/A | 22.15 | 41.4 |
| SF10P | Cs5 | *CsFT3* | 8 | 7038258 | 8.8E-10 | 8 | G/A | 17.53 | 24.6 |
|  | Cs2 | *CsCFL1* | 9 | 30838575 | 7.8E-09 | 5 | C/A | 18.87 | 41.3 |
|  | Cs10 | - | X | 85663484 | 1.8E-08 | 16 | G/T | -8.66 | 10.8 |
| FT50I | Cs9 | - | 7 | 7258159 | 5.2E-31 | 26 | A/- | 24.32 | 30.6 |
|  | Cs5 | *CsFT3* | 8 | 7038258 | 8.2E-10 | 8 | G/A | 15.49 | 20.9 |
|  | Cs2 | *CsCFL1* | 9 | 30838575 | 3.5E-13 | 5 | C/A | 23.79 | 24.3 |

<sup>a</sup> Complete name of key introduced loci: *AutoFlower3* (*CsFT3*); *CircadianFloweringLocus1* (*CsCFL1*); *FloweringTime\_Maturity4* (*CsFT4*).
<sup>b</sup> Genomic position of the significant SNP in base pairs, reported based on the *Cannabis* reference genome cs10 v2 (GenBank accession no. *GCA\_900626175.2*).
<sup>c</sup> Minor Allele Frequency.
<sup>d</sup> Major/minor.
<sup>e</sup> Estimated allelic effect of the minor allele on the trait.
<sup>f</sup> Proportion of Phenotypic Variance Explained by the significant SNP.

#### 3.3.2 Morphological characteristics

A total of 52 significant genetic markers were identified across 20 morpho-agronomic traits in the GWAS panel. The results for these traits are grouped into three categories: (i) Node and Branching Architecture, (ii) Growth and Structural Dimensions, and (iii) Biomass Yield.

##### i. Node and Branching Architecture

In the Node and Branching Architecture subcategory, 11 significant genetic markers were associated with five traits, demonstrating phenotypic contributions with PVE values spanning from 3.7% (Cs18 associated with NTFIS) up to an impressive 87.1% (Cs16 associated with NMI) (Fig. 5a; Table 2; Fig. S14b). Key loci identified include *NodeNumberRegulator* (*CsNNR1*), represented by marker Cs11 (Chr3:90476582; PVE: 25.2% for NNH and 74.2% for NTFIS), *CsNNR2* (Cs19 on Chr1:12025015; PVE: 51.4% for NTMI), *TopNodesDensity* (*CsTND*), which includes two complementary loci: *CsTND1* (Cs12 on Chr4: 79659045; PVE: 23.6% for NNH) and *CsTND2* (Cs16 on Chr4:79296939; PVE: 87.1% for NMI) as well as *CsCFL1* (Cs2 on Chr9:30838575; PVE: 47% for NLS). The *CsFT3* locus, previously represented in flowering time traits by Cs1 and Cs5, is characterized by Cs5 (Chr8:7038258; PVE: 7.8% for NLS) and Cs13 (Chr8:7035841; PVE: 35% for NNH and 34.3% for NTMI), highlighting its extended influence on plant architecture. The allelic impact of Cs16 (*CsTND2*) was particularly pronounced, where individuals carrying the ‘CC’ allele exhibited a significantly higher node count on the main inflorescence (12.23 nodes) compared to those with the ‘AA’ allele (6.8 nodes) (Fig. S16). Together, these loci coordinate both general and region-specific control over phyllotactic development, including basal-to-apical node and branch patterning.

**Fig. 5.**
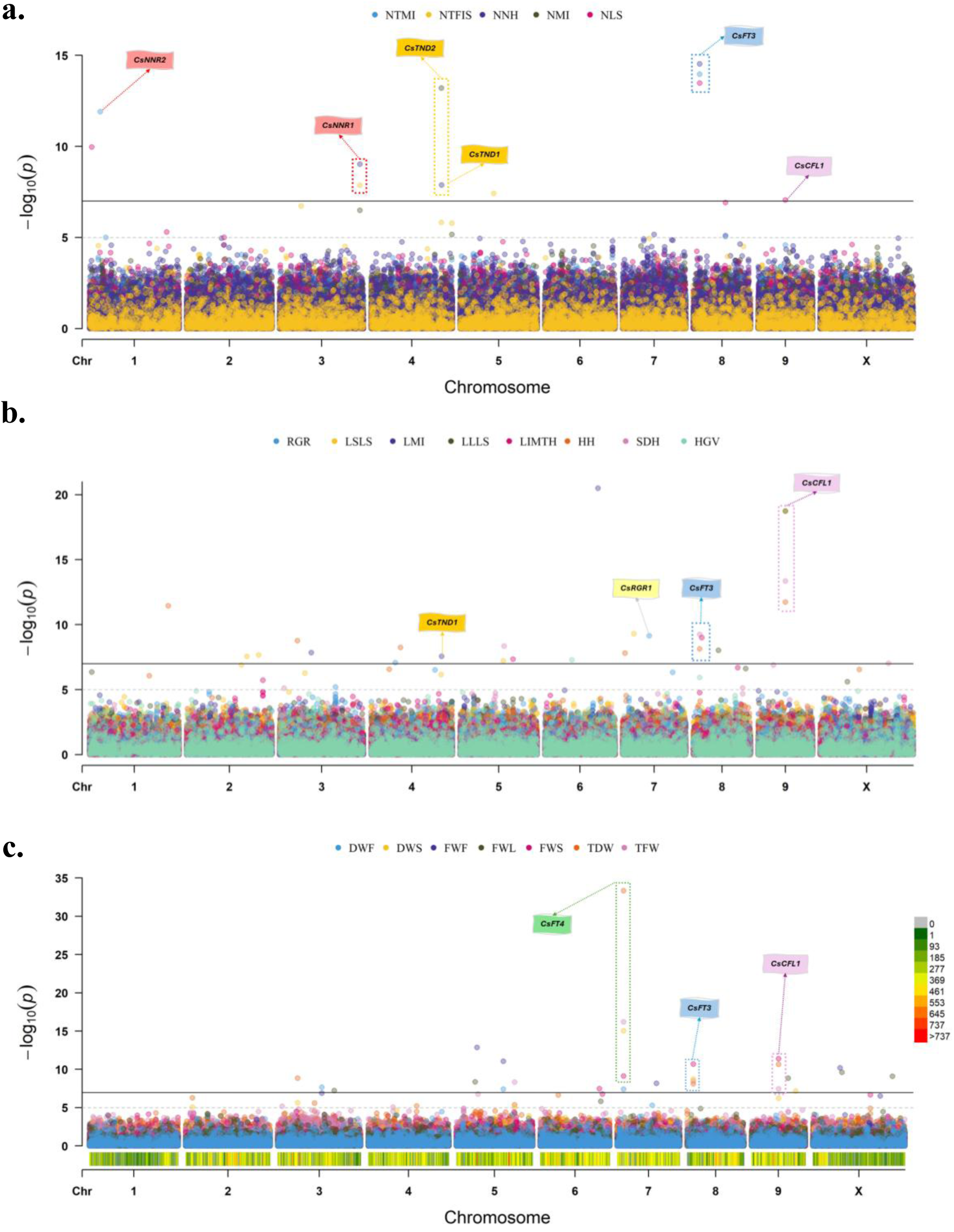
Manhattan plots illustrating GWAS results for morpho-agronomic traits. Plots show associations for Node and Branching Architecture traits (**a**), Growth and Structural Dimension traits (**b**), and Biomass Yield traits (**c**).

**Table 2.**
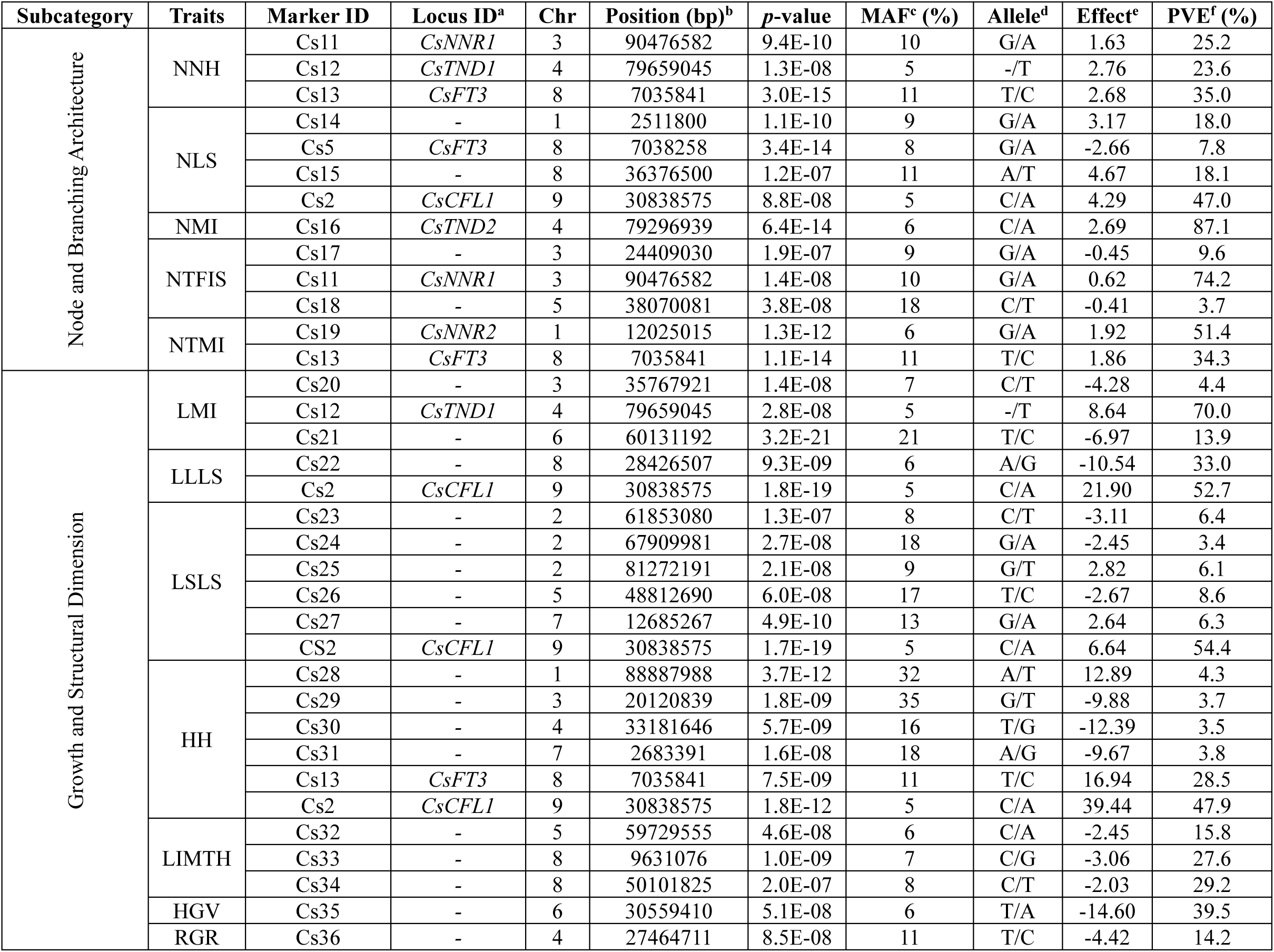
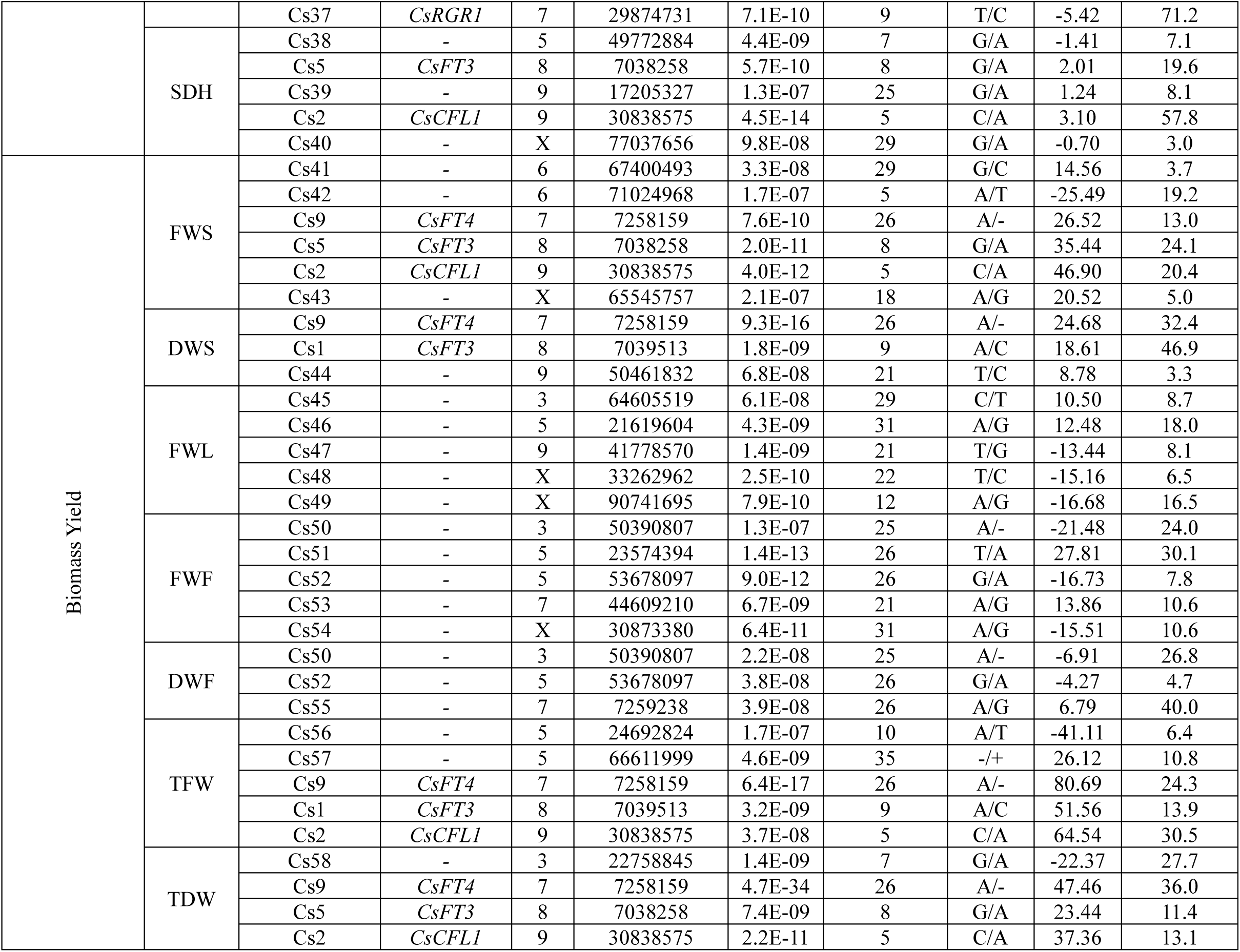
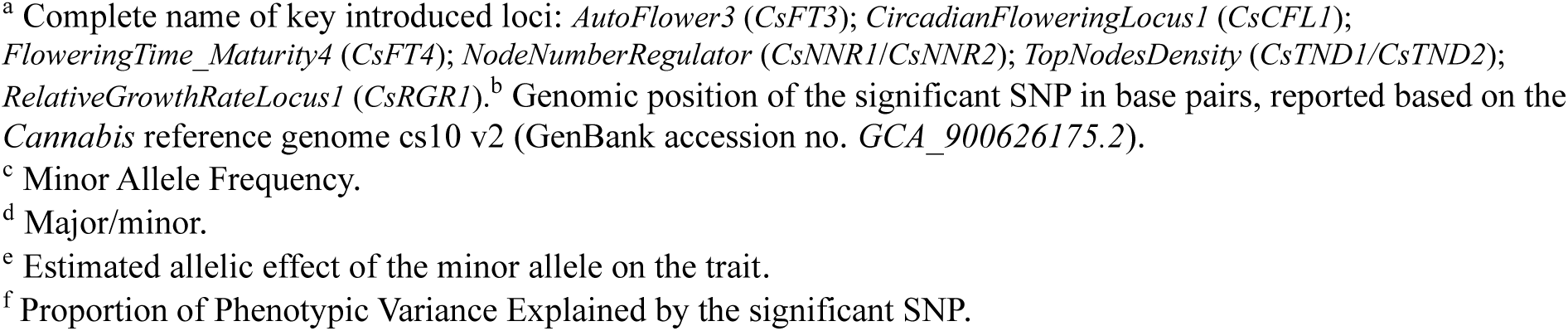
List of 52 markers associated with 20 morpho-agronomic traits in the cannabis landrace panel.

##### ii. Growth and Structural Dimension

GWAS analysis identified 25 significant markers associated with eight Growth and Structural Dimension traits, with phenotypic effects spanning 3% (Cs40 for SDH) to 71.2% (Cs37 for RGR) (Fig. 5b; Table 2; Fig. S14c). Notable loci included *RelativeGrowthRateLocus1* (*CsRGR1*), represented by marker Cs37 (Chr7:29874731; PVE: 71.2% for RGR), which demonstrated the strongest individual marker effect in this trait category, closely followed by *CsTND1* (Cs12 on Chr4:79659045; PVE: 70% for LMI). Together, *CsTND1* and *CsTND2* underline a coordinated genetic control over inflorescence architecture, through the modulation of both node number and internodal length. Additionally, *CsCFL1* (Cs2 on Chr9: 30838575) showed substantial pleiotropic effects with PVE values of 52.7% for LLLS, 54.4% for LSLS, 47.9% for HH and 57.8% for SDH. Additionally, *CsFT3* locus was represented by Cs5 (Chr8:7038258; PVE: 19.6% for SDH) and Cs13 (Chr8:7035841; PVE: 28.5% for HH), further supporting its role in plant architecture beyond flowering regulation. Several markers associated with different growth and structural dimension traits were located in close proximity, suggesting shared genomic regions influencing these correlated characteristics. The allelic impact of these loci was particularly pronounced. For *CsRGR1*, individuals carrying the ‘CC’ allele exhibited a significantly higher RGR (104.17 mg.g^-1^.day^-1^) compared to those with the ‘TT’ allele (69.7 mg.g^-1^.day^-1^). Similarly, for *CsCFL1*, the ‘CC’ allele consistently promoted increased structural dimensions across multiple traits (HH: 161.38 vs 85.29 cm; LLLS: 53.72 vs 28.4 cm; LSLS: 14.87 vs 6.85 cm; SDH: 13.6 vs 7.67 mm), demonstrating its broad architectural influence compared to the ‘AA’ allele. Overall, these markers coordinate fundamental aspects of plant morphology and growth dynamics, offering valuable targets for architectural improvement in breeding programs (Fig. S17).

##### iii. Biomass Yield

Biomass yield analysis identified 22 significant markers spanning seven yield components, with effects ranging from 3.3% (Cs44 for DWS) to 46.9% (Cs1 for DWS) (Fig. 5c; Table 2; Fig. S14d). The genetic architecture was largely shaped by the *CsFT3* region, where marker Cs1 (Chr8:7039513) exerted the strongest effect, particularly on stem dry weight (DWS; PVE = 46.9%) and also contributed to total fresh weight (TFW; 13.9%). Meanwhile, its closely linked counterpart, Cs5 (Chr8:7038258), displayed a partially complementary effect pattern, with major influence on stem fresh weight (FWS; PVE = 24.1%) alongside a notable contribution to total dry weight (TDW; 11.4%). The *CsFT4* locus emerged as a major yield regulator through Cs9 (Chr7:7258159), controlling multiple biomass components with remarkable consistency: 32.4% PVE for DWS, 36.0% for TDW, 24.3% for TFW and 13% for FWS. Additionally, *CsCFL1* (Cs2 on Chr9:30838575; PVE: 20.4% for FWS, 30.5% for TFW and 13.1% for TDW) extended its pleiotropic influence into yield determination through enhanced fresh biomass accumulation and total plant weight. The *CsFT4* locus demonstrated exceptional yield enhancement, where presence of the ‘AA’ allele nearly doubled TDW (114.92 g vs 64 g) and increased TFW by 79% (276.47 g vs 154.41 g) compared to the deletion variant. The *CsFT3* region also showed similarly dramatic effects, with the ‘AA’ allele of Cs1 producing a six-fold higher DWS (51.73 g vs 8.41 g) compared to the ‘CC’ allele. Consistent with its broad developmental control, the *CsCFL1* locus (Cs2) with the ‘CC’ allele showed substantial 2.7-fold gain in TDW (94.94 g vs 35.58 g), a 2.5-fold increase in TFW (229.81 g vs 91.87 g) and a 3.3-fold gain in FWS (112.3 g vs 34.16 g) compared to the heterozygous ‘AC’ allele (Fig. S18). It is noteworthy that while the DWL trait was included in the GWAS for biomass yield, no significant markers were identified for this specific trait. These yield-controlling loci represent prime candidates for marker-assisted selection targeting biomass optimization.

#### 3.3.3 Cannabinoid Profiles

GWAS of cannabinoid profiles identified 34 significant genetic markers associated with 14 phytochemical (cannabinoid) traits. The results for these traits are categorized by sub-group.

##### i. THC-related

THC-related traits analysis identified 20 significant markers associated with nine major biochemical traits, with PVE ranging from 3% (Cs69 for CBC) to 94% (Cs59 for Total THC Potential) (Fig. 6a; Table 3; Fig. S14e). The cannabinoid biosynthetic pathway was primarily regulated by *THCALocus1* (*CsTHCAL1*), where the marker Cs59 (Chr5:29925798) exerted dominant effects across multiple THC-related metabolites, explaining 94% of the variance in Total THC Potential, 91.4% in THCA, 89.9% in CBN, and 72.9% in Δ^9^-THC. Beyond *CsTHCAL1*, other major loci shaped specific branches of the biosynthetic network: *CBGALocus1* (*CsCBGAL1*) was captured by marker Cs67 (Chr3:13775640; PVE: 90.6% for CBGA and 7% for CBG), *CBGRegulator1* (*CsCBGR1*) by Cs68 (Chr5:24473236; PVE: 91.3% for CBG), and THC:*CBDRatioLocus1* (*CsTCBR1*) by Cs66 (Chr1:24225734; PVE: 79.2% for THC:CBD ratio), highlighting distinct control over precursor and end-product partitioning. The THCV biosynthesis appeared to be regulated by Cs64 (Chr5:2418037; PVE: 57.9%), designated as *THCVRegulator1* (*CsTHCVR1*). In contrast to these high-effect loci, CBC biosynthesis was governed by a more complex multi-locus architecture, involving 10 markers across chromosomes 1, 3, 4, 6, 9, and X, with the strongest individual contribution from Cs78 (ChrX:48786650; PVE: 27.6%), suggesting a polygenic basis for this trait.

**Fig. 6.**
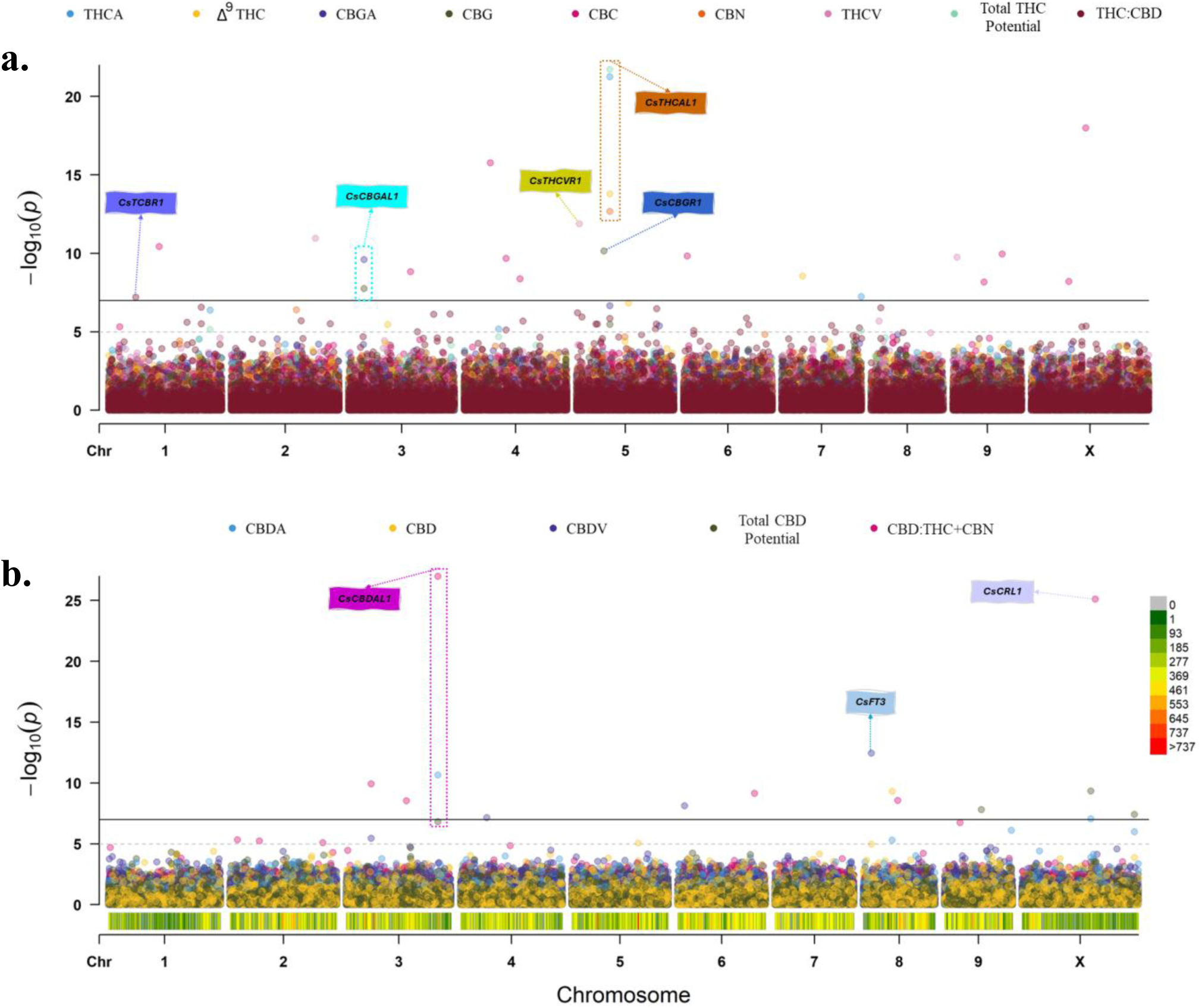
Manhattan plots illustrating GWAS results for cannabinoid traits. Plots show associations for THC-related traits (**a**) and CBD-related traits (**b**).

**Table 3.**
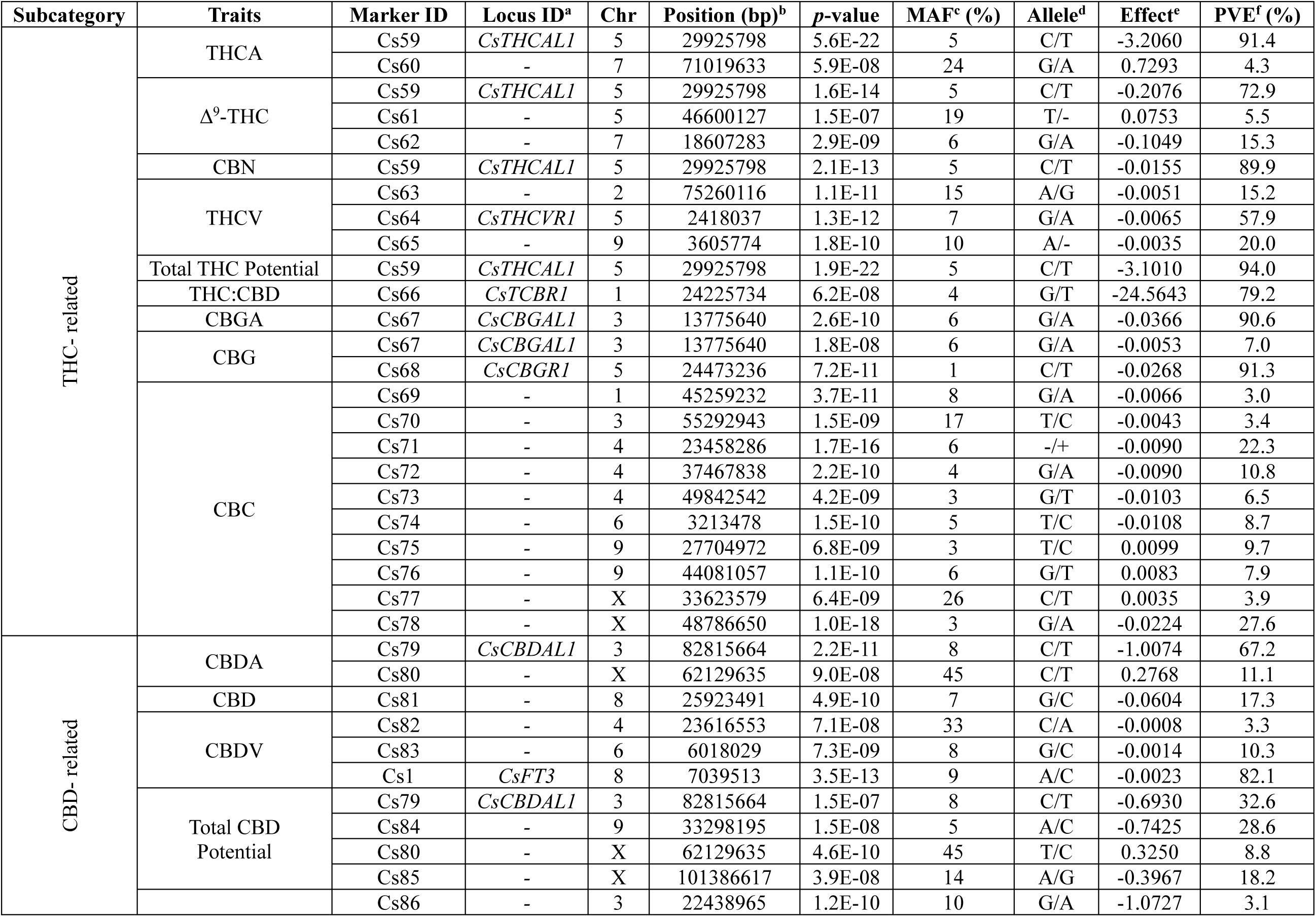
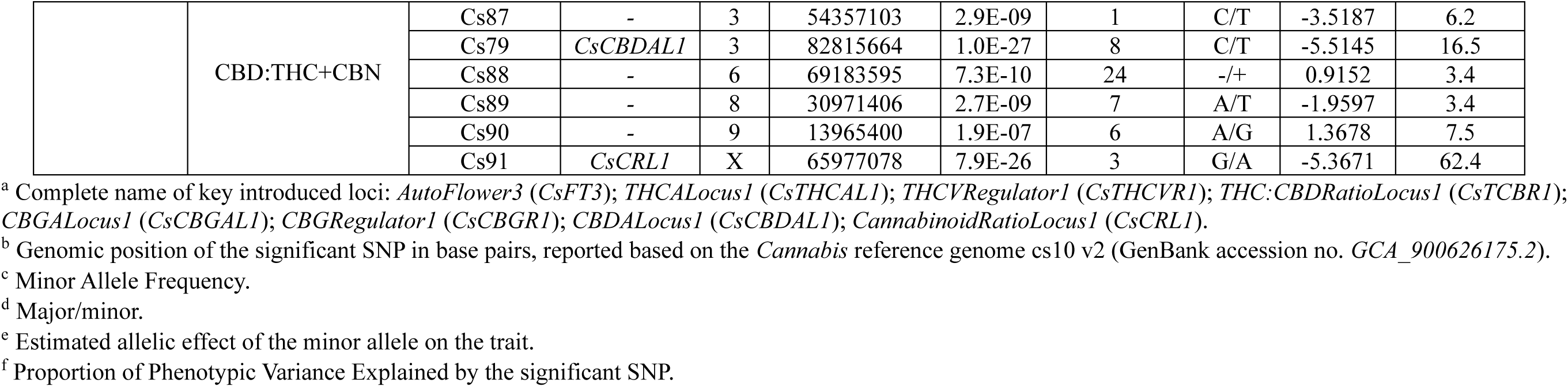
List of 34 markers associated with 14 phytochemical (cannabinoid) traits.

The *CsTHCAL1* locus demonstrated exceptional potency enhancement, where the ‘TT’ allele produced nearly six-fold higher Total THC Potential (9.4% vs 1.65% in ‘CC’ allele) and similar magnifications in THCA content (9.84% vs 1.66%). For the precursor pathway, *CsCBGAL1* showed the ‘AA’ allele producing substantially elevated CBGA levels (0.178% vs 0.04% in ‘GG’ genotypes), representing a 4.5-fold increase in this critical biosynthetic intermediate. The *CsTCBR1* exhibited striking chemotype differentiation. The ‘TT’ allele shifting THC:CBD ratios from 7.32 in ‘GG’ backgrounds to 59.73 in ‘TT’, indicating an 8-fold change in cannabinoid profile direction Fig. S19.

##### ii. CBD-related

CBD-related traits through GWAS revealed 14 significant markers controlling five CBD-related traits, with genetic contributions spanning 3.1% (Cs86 for CBD:THC+CBN) to 82.1% (Cs1 (*CsFT3*) for CBDV) (Fig. 6b; Table 3; Fig. S14f). The *CBDALocus1* (*CsCBDAL1*), anchored by Cs79 (Chr3:82815664), emerged as a pivotal regulator with the highest individual effect on CBDA production (67.2% PVE) and substantial influence on CBD:THC+CBN ratios (16.5% PVE) and Total CBD Potential (32.6% PVE). The X-linked *CannabinoidRatioLocus1* (*CsCRL1*), represented by Cs91 (ChrX:65977078), demonstrated exceptional control over cannabinoid ratios with 62.4% PVE for CBD:THC+CBN, establishing sex-linked inheritance patterns in cannabinoid composition. Remarkably, the flowering-time regulator *CsFT3* extended its pleiotropic influence into cannabinoid metabolism, where Cs1 (Chr8:7039513) achieved the strongest single marker effect in this category (82.1% PVE for CBDV). For *CsCBDAL1*, the ‘TT’ allele dramatically enhanced CBDA production (2.69% vs 1.11% w/w in ‘CC’) and substantially increased CBD:THC+CBN ratios (7.9 vs 1.09 in ‘CC’), establishing its role as a major cannabinoid biosynthesis enhancer. Most remarkably, *CsFT3* revealed an unexpected connection between flowering regulation and cannabinoid metabolism, where the ‘CC’ allele produced eight-fold higher CBDV concentrations (0.0097% vs 0.0012% w/w in ‘AA’), highlighting novel pleiotropic pathways linking developmental timing to secondary metabolite production (Fig. S20).

### 3.4 Haplotype block and pleiotropic loci characterization

To understand the genetic basis of observed phenotypic variation, candidate genes within HB regions encompassing 91 unique genome-wide significant markers associated with 40 morphological, phenological, and cannabinoid traits were investigated (Table S14). These HBs, defined by variants with an r^2^ ≥ 0.75, varied considerably in size, ranging from 5 bp (e.g., Cs7) to 37.6 kb (e.g., Cs45). Notably, several significant markers were found within the same haploblock, such as Cs1, Cs5, and Cs13 on chromosome 8 (HB size: 3 kb). This analysis identified 34 putative candidate genes within these regions, although six were classified as having uncharacterized or unknown functions, providing clear avenues for future investigation (Table S14).

A key finding of this study is the extensive pleiotropy observed for 15 major loci, which are represented by 17 key markers (Fig. 7). These loci exert broad regulatory control across multiple, seemingly distinct trait categories, highlighting genomic hotspots with far-reaching effects. Specifically, the *CsFT3* complex, located within the 3 kb haploblock on chromosome 8, serves as a prime example of pleiotropy. The markers within this locus (Cs1, Cs5, and Cs13) significantly influenced multiple phenological stages (GVP, SFFI, SFFP, SF10I, SF10P, FT50I), morphological traits (NNH, NTMI, HH, NLS, SDH), and biomass traits (FWS, TDW, DWS, TFW), with marker Cs1 also showing a strong association with CBDV concentration (82.1% PVE). Furthermore, marker Cs2 (*CsCFL1*) on chromosome 9 displayed the broadest pleiotropic impact, associated with 12 distinct traits across phenological, morphological, and biomass categories, and its alleles contribute to the differentiation of early, medium, and late-flowering accessions. *CsFT4* (Cs9) on chromosome 7 was identified as a homolog of *CsFT3*, influencing later phenological stages and several biomass components. Similarly, the loci *CsNNR1* (Cs11) on chromosome 3 and its homolog *CsNNR2* (Cs19) on chromosome 1 were found to regulate node number and branching patterns. The loci *CsTND1* (Cs12) and *CsTND2* (Cs16) on chromosome 4 demonstrated specialized pleiotropy, with *CsTND1* controlling node number and internode length and *CsTND2* specifically regulating the number of nodes on the main inflorescence, thereby affecting main inflorescence size and density. The extensive pleiotropy observed for several key markers emphasizes the complexity of trait regulation and offer valuable resources for molecular breeding strategies in cannabis.

**Fig. 7.**
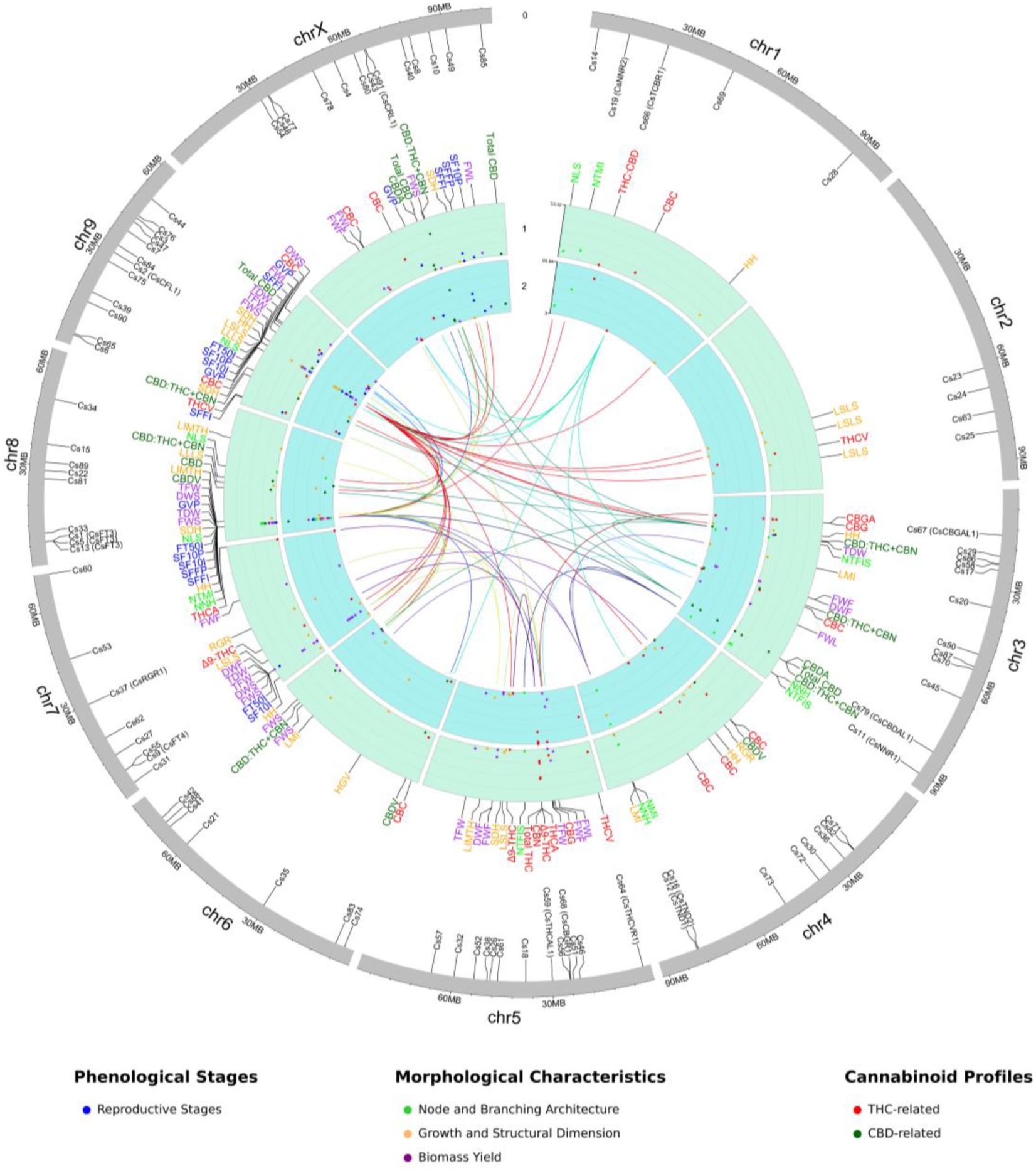
Circular plot illustrates the genome-wide distribution of 91 significant markers and their associated 40 traits identified by GWAS. The outermost ring (0) displays chromosomes (1 to X) and the position of the identified markers. The first inner ring (1) represents the transformed *p*-values (−log_10_ (*p*)) from GWAS, while the second inner ring (2) shows the proportion of phenotypic variance explained (PVE). Labels on these inner rings correspond to the various traits, positioned on the first inner ring. The different colors within rings 1 and 2, along with the trait labels, indicate distinct phenological, morphological, and phytochemical trait categories. Connecting lines in the center of the plot link markers associated with multiple distinct traits across the genome, highlighting pleiotropic effects and genomic hotspots.

## 4. Discussion

This study provides a comprehensive genetic and phenotypic characterization of cannabis landrace accessions, comprising both females and males, offering valuable insights into the genetic architecture underlying key phenological, morphological, and phytochemical traits. Utilizing high-density GBS data (Torkamaneh et al. 2021) and GWAS, we identified 91 unique genome-wide significant markers associated with 40 traits, providing valuable insights into genetic architecture underlying trait variation in this crop. Our most significant finding is the discovery of 15 highly pleiotropic loci with exceptional effects on multiple trait categories, particularly *CsFT3* and *CsCFL1*, which demonstrate that single genomic regions can simultaneously control flowering time, plant architecture, biomass production, and cannabinoid biosynthesis. Notably, *CsTHCAL1* and *CsCBDAL1* exhibited exceptionally high phenotypic variance explained (94% for multiple THC-related and 67.2% for CBDA, respectively), underscoring their relevance in regulating cannabinoid pathways. These discoveries provide unprecedented insights into the genetic mechanisms governing cannabis diversity and offer powerful tools for molecular breeding programs.

The remarkable allelic diversity uncovered in our cannabis landrace panel mirrors the transformative role landraces have played in other crops (e.g., maize, rice, and wheat) through the identification of key adaptive alleles (Zhang 1995; Rubiales and Niks 2000; Ren et al. 2005; Hurni et al. 2015; Dias et al. 2024). Our Iranian cannabis landraces provided the genetic diversity essential for identifying causal variants that remain hidden in narrow commercial germplasm (de Ronne et al. 2024; de Ronne and Torkamaneh 2025). LD analysis revealed a notably rapid decay rate, much faster than in previous cannabis studies, which range from 3.9 kb (Basal) to 6 kb (Drug-type) (Ren et al. 2021), 22.6-89 kb in commercial Canadian drug-type cannabis cultivars (de Ronne et al. 2024) and 6.7 kb in feral germplasm from the United States (Aina et al. 2025). This aligns with outcrossing or self-incompatible species such as tea (100 bp–8 kb; Lei et al., 2023) and maize (500 bp–6.3 kb; Dinesh et al., 2016; Pavan et al., 2020; Remington et al., 2001; Yan et al., 2009), and contrasts with self-pollinated crops like soybean (>100 kb), tomato (1 Mb), rice (150 kb), and wheat (8 Mb) (Liu et al. 2017a, b, 2019; Torkamaneh et al. 2020a; Viana et al. 2022). This rapid LD decay, shaped by dioecy, wind pollination, and limited selection pressure, enables high-resolution mapping and demands dense marker coverage (Mohammadi et al. 2020; Belzile and Torkamaneh 2022; Hyten 2022). Interestingly, within the subpopulations, the highest diversity value in our panel was nearly identical to the lowest value observed among subpopulations of commercial drug-type genotypes (8.44 ×10^-4^) (de Ronne et al. 2024). This contrasts sharply with broader populations, where *θ_π_* values can reach as high as 3.87×10^-3^, reflecting much greater genetic diversity (Ren et al. 2021). Consistent with our previous study (Babaei et al. 2024), this population also showed substantial phenotypic variation across key traits. Heritability estimates (H² and SNP-based h²) revealed strong additive genetic control, supporting high selection potential (Falconer and Mackay 1996; Wray and Visscher 2008; Naim-Feil et al. 2021). For example, H² for RGR reached 0.95 in females and 0.99 in males, with striking sex-specific differences in morphological traits like DWF (0.33 vs. 0.87) and FWF (0.23 vs. 0.92) (Babaei et al. 2024). The observed deviations from normal distribution, indicative of rare alleles contributing to extreme phenotypes, further emphasize the utility of landraces in capturing unique genetic variations often absent in elite breeding lines (Zhu et al. 2008; Ingvarsson and Street 2011; Gupta et al. 2019; Mohammadi et al. 2020; Torkamaneh et al. 2020a; Belzile and Torkamaneh 2022).

Comparing our HD-GBS results with de Ronne et al. (2024) on commercial Canadian drug-type cannabis shows similar SNP counts, densities, and gene-annotated SNP proportions. However, the commercial genotypes exhibited higher MAF (21.7%) and heterozygosity (25.5%) due to intensive selective breeding and use of diverse parental lines, which stabilize traits and maintain allele frequencies. In contrast, landraces, shaped by prolonged natural selection and limited human intervention, exhibit lower MAF reflecting broader allelic diversity and rare alleles at low frequencies (Linck and Battey 2019; Kovalchuk et al. 2020). Studies by Sawler et al. (2015) and Soler et al. (2017) confirm hemp populations have higher heterozygosity (16% and 40.5%) than drug-types (12.5% and 28.2%), likely due to wider genetic bases and hybridization levels. Lynch et al. (2016) reported lower heterozygosity in European hemp (22%) vs. drug-types (31%), while Gao et al. (2014) found higher heterozygosity in Chinese hemp (35.5–37%) compared to European hemp (18.2%), highlighting geographic and breeding history effects. Overall, these patterns reflect natural selection and limited hybridization in landraces versus targeted breeding in commercial cannabis to stabilize specific traits (Hurgobin et al. 2021). Thus, the extensive phenotypic and genetic diversity, coupled with strong heritability, and rapid LD decay provides a robust foundation for GWAS and targeted breeding (Ingvarsson and Street 2011; Alqudah et al. 2020; Belzile and Torkamaneh 2022; Torkamaneh and Belzile 2022).

The genetic architecture of Iranian cannabis landraces, characterized by three distinct genetic clades, reflects the ecological and historical complexity of Iranian Plateau. Its fragmented topography and climatic heterogeneity have fostered local adaptation, creating recombination hotspots and divergent subpopulations (Akhani et al. 2010; Djamali et al. 2012; Shumilovskikh et al. 2016; Manafzadeh et al. 2017; Gurjazkaite et al. 2018). Iran’s role as a Bronze Age trade hub, notably along the Silk Road, likely facilitating the dispersal and hybridization of cannabis (Warf 2014; McPartland 2018; Kovalchuk et al. 2020; Abdullaev 2022; Liu and Brancaccio 2022; Ndlangamandla et al. 2024).

While Ren et al. (2021), proposed a single East Asian origin based on Chinese hemp, earlier hypotheses favored Central Asia (de Candolle 1867; De Candolle 1883; McPartland 2017, 2018; McPartland and Small 2020). Notably, Ren’s study excluded Iranian accessions, potentially missing key diversification events. Supporting this, Balant et al. (2025) identified Iranian samples as a genetically distinct subgroup, favoring a geography-based classification over use-type model (Ren et al. 2021). Our genomic analysis revealed three Iranian subpopulations shaped by geography, environment, and traditional selection (Ren et al. 2021). Similar to Chinese cannabis (Chen et al. 2022), Iranian landraces show ecotype-based adaptation. Clade I’s intermediate phylogenetic position suggests either ancestral ties to Clades II and III or historical hybridization (Ramos-Madrigal et al. 2019; Pérez-Escobar et al. 2021, 2022). Clade III stands out genetically, nearing thresholds used in the Genetic Species Concept (Bradley and Baker (2001), though such metrics remain debated (Zachos 2016). Despite phenotypic variation—from ruderalis-like too tall, late-flowering types—Clade III forms a cohesive genetic group, consistently CBD-dominant. This highlights evolutionary processes like hybridization and convergent evolution that blur species boundaries (Steenwyk et al. 2023).

The observed phenotypic diversity in cannabis is profoundly influenced by environmental factors, including geographical latitude, climate, and the selective pressures exerted by both natural and human forces (Hazekamp and Fischedick 2012; Babaei and Ajdanian 2020; Ren et al. 2021; Babaei et al. 2022, 2025; Lata et al. 2023). As a short-day plant, cannabis synchronizes its biological activities, including the critical transition to flowering, with environmental rhythms through its internal circadian clock. This fundamental biological process acts as an upstream regulator, influencing not only flowering time but also cascading effects on plant architecture, growth, and biomass accumulation (Webb 2003; Harmer 2010; Gottlieb 2019; Webb et al. 2019; Steed et al. 2021). For instance, key loci identified in this study, such as *AutoFlower3* (*CsFT3*) on chromosome 8 (encompassing markers Cs1, Cs5, and Cs13) and *CircadianFloweringLocus1* (*CsCFL1*) (marker Cs2 on chromosome 9), exemplify these broad regulatory impacts. These markers exhibit significant pleiotropic effects, influencing multiple traits across phenological and morphological (e.g., node and branching architecture, growth dimensions), and cannabinoid profiles, consistent with their roles in photoperiod and circadian rhythm regulation. Such pleiotropy, where a single genomic region affects multiple seemingly unrelated traits, is a common feature in plant development and adaptation, observed in other short-day plants like soybean and rice (e.g., *FLOWERING LOCUS T* (*FT*) homologs influencing vegetative and reproductive branching and yield-related traits), and long-day plants like Arabidopsis (e.g., *FLOWERING LOCUS T* (*FT*), *FRIGIDA* (*FRI*), *FLOWERING LOCUS C* (*FLC*) affecting branching, seed germination, and water use efficiency) (Hiraoka et al. 2013; Cao et al. 2022; Weng et al. 2022; Maple et al. 2024; Mohamedikbal et al. 2024). The identification of *CsFT3* and *CsCFL1* in our study, with their broad pleiotropic effects, strongly suggests their central role in this intricate regulatory network in cannabis landraces. We also identified specific loci with major effects on phytochemical traits. For instance, *CsTHCAL1* on chromosome 5 and *CsCBDAL1* on chromosome 3 are directly linked to the primary biosynthesis pathways of THC and CBD, respectively. The genetic architecture of THC and CBD extends beyond canonical synthase genes, with contributing markers distributed across various chromosomes, and total cannabinoid content QTL often residing outside the synthase cluster (Hurgobin et al. 2021; de Ronne and Torkamaneh 2025). Our phenotypic analysis, which categorized cannabinoids into THC-related and CBD-related groups based on strong correlations and PCA, aligns with their distinct biosynthetic pathways (Govindarajan et al., 2023; Gülck and Møller, 2020; Hurgobin et al., 2021; Ingvardsen and Brinch-Pedersen, 2023; Welling et al., 2020).

## 5. Conclusion

In summary, this study provides a robust genetic framework for understanding trait variation in *Cannabis* landraces, identifying key markers linked to important traits. These findings offer valuable tools for marker-assisted and gene-editing approaches, supporting the development of improved cultivars. Preserving these landraces is crucial for maintaining genetic diversity and unlocking novel traits for future breeding innovations.

## Abbreviations

BIC: Bayesian Information Criterion
BLINK: Bayesian-information and linkage-disequilibrium iteratively nested keyway
CBC: Cannabichromene
CBD: Cannabidiol
CBDA: Cannabidiolic acid
CBDV: Cannabidivarin
CBG: Cannabigerol
CBGA: Cannabigerolic acid
CBN: Cannabinol
*CsCBDAL1*: *CBDALocus1*
*CsCBGAL1*: *CBGALocus1*
*CsCBGR1*: *CBGRegulator1*
*CsCFL1*: *CircadianFloweringLocus1*
*CsCRL1*: *CannabinoidRatioLocus1*
*CsFT3*: *AutoFlower3*
*CsFT4*: *FloweringTime_Maturity4*
*CsNNR1*: *NodeNumberRegulator1*
*CsNNR2*: *NodeNumberRegulator2*
*CsRGR1*: *RelativeGrowthRateLocus1*
*CsTCBR1*: *THC:CBDRatioLocus1*
*CsTHCAL1*: *THCALocus1*
*CsTHCVR1*: *THCVRegulator1*
*CsTND1*: *TopNodesDensity1*
*CsTND2*: *TopNodesDensity2*
CV: Coefficient of Variation
DAPC: Discriminant Analysis of Principal Components
DAS: Days after sowing
DWF: Dry Weight of Flowers
DWL: Dry weight of leaves
DWS: Dry weight of stems
FDR: False Discovery Rate
FT50I: Flowering Time 50% in Individuals
FWF: Fresh Weight of Flowers
FWL: Fresh weight of leaves
FWS: Fresh weight of stems
GBS: Genotyping-by-sequencing
GO: Gene Ontology
GVP: GV Point
GWAS: Genome-wide association studies
H^2^: Broad-sense heritability
h^2^: SNP-based heritability
HB: Haplotype Block
HH: Height on Harvest day
HGV: Height to GV Point
LD: Linkage Disequilibrium
LIMTH: Length of Internode in the Middle Third of the main stem on Harvest day
LLLS: Length of Longest Lateral Shoot
LMI: Length of Main Inflorescence
LOD: Limit of Detection
LSLS: Length of Shortest Lateral Shoot
MAF: Minor Allele Frequency
MAS: Marker-assisted selection
MNPs: Multi-nucleotide polymorphisms
MQ: Mapping Quality
MSE: Mean Squared Error
NGS: Next-generation sequencing
NMI: Number of Nodes on the Main Inflorescence
NNH: Number of Nodes on the main stem on Harvest day
NLS: Number of Lateral Shoot
NRMSE: Normalized Root Mean Squared Error
NTFIS: Number of Nodes to the First Lateral Shoot
NTMI: Number of Nodes to the Main Inflorescence
PCA: Principal Component Analysis
PCs: Principal components
PVE: Proportion of Phenotypic Variance Explained
QUAL: Quality
QTL: Quantitative Trait Loci
RCBD: Randomized complete block design
RGR: Relative Growth Rate
RMSE: Root Mean Square Error
SDH: Stem Diameter on Harvest day
SF10I: Start 10% Flowering Time in Individuals
SF10P: Start 10% Flowering Time in 50% Population
SFFI: Start Flower Formation Time in Individuals
SFFP: Start Flower Formation Time in 50% Population
SNP: Single nucleotide polymorphism
SVs: Structural variations
TDW: Total Dry Weight
TFW: Total Fresh Weight
THC: Δ^9^-tetrahydrocannabinol
THCA: Δ^9^-tetrahydrocannabinolic acid
THCV: Tetrahydrocannabivarin

## Author Contributions

M.B: Conceptualization; methodology; investigation; data curation; formal analysis; resources; visualization; writing – original draft; writing – review & editing. D.T: Conceptualization; supervision; funding acquisition; resources; writing – review & editing.

## Data Availability

The raw datasets generated during the current study are available. Processed data supporting the findings are publicly accessible via Figshare (DOI: <u>10.6084/m9.figshare.29868215</u>).

## Ethics approval

Seed collection and cultivation were conducted with ethics approval and license from the anti-narcotics police in Razavi Khorasan. Activities at Université Laval complied fully with Health Canada regulations under license LIC-QX0ZJC7SIP-2021.

## Key Message

Our GWAS of *Cannabis* landraces uncovered 15 pleiotropic loci and 91 trait-associated regions, offering a rich genomic resource for precision breeding and cultivar improvement.

## Acknowledgements

The authors wish to thank Ladan Ajdanian and Perrine Feutry for their valuable contribution in data collection.

