## Supplemental Figures S1 to S20 for "Genetic architecture of phenological, morphological, and phytochemical traits in *Cannabis* landraces"

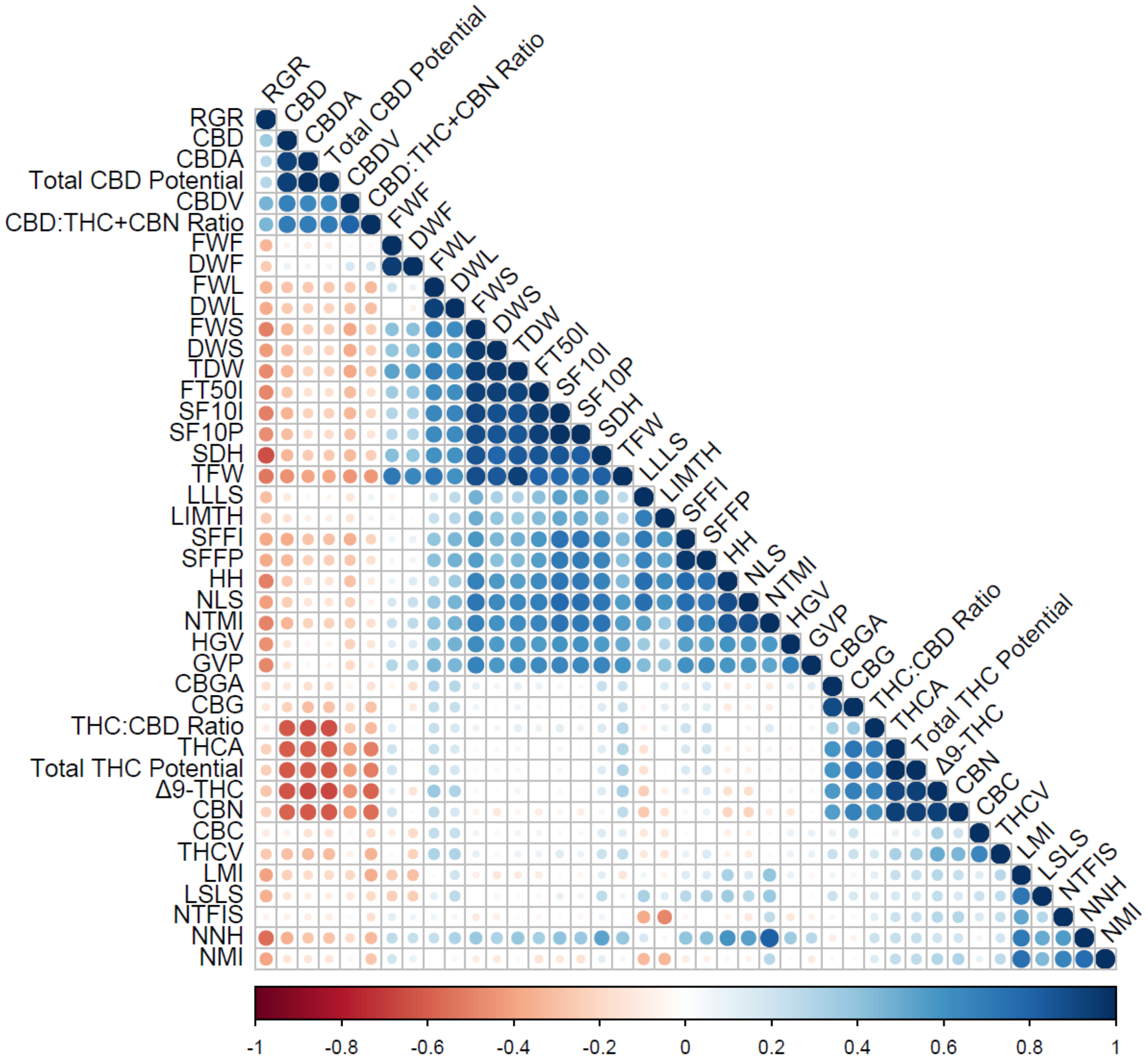


**Figure S1:** Pairwise Pearson's correlation coefficients among 41 morphological, phenological, and cannabinoid traits in 145 *Cannabis* landrace accessions. The correlogram illustrates the strength and direction of linear relationships between evaluated traits. Blue circles indicate positive correlations, while red circles indicate negative correlations. The intensity of the color and the size of the circles are proportional to the absolute value of the correlation coefficient. Non-significant correlations (P>0.05) are left blank.

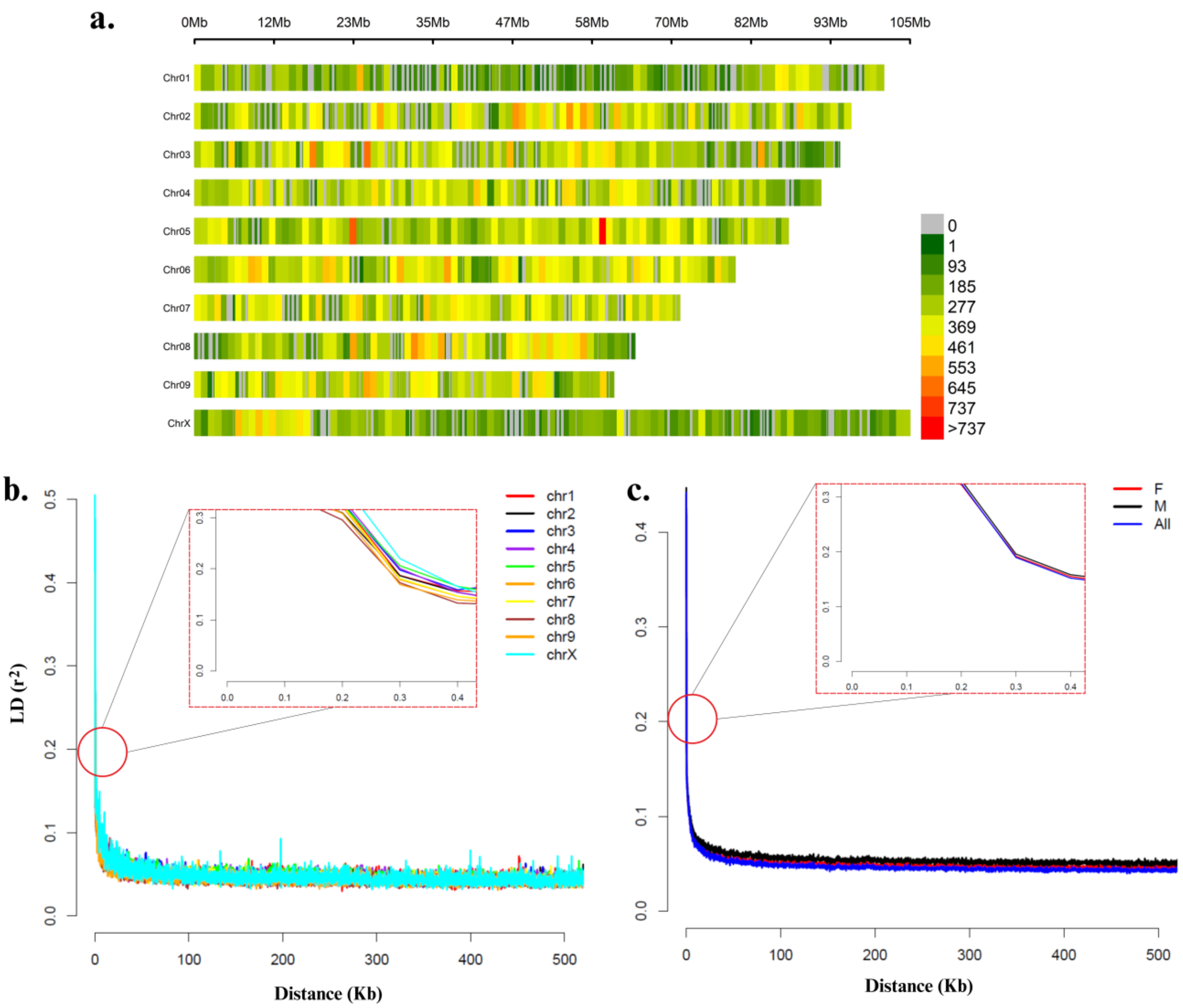


**Figure S2:** Genome-wide SNP distribution and linkage disequilibrium (LD) in 145 *Cannabis* landrace accessions. (**a**) Chromosomal SNP density across the genome, visualized using a 1 Mb window, highlighting variations in marker distribution among chromosomes. (**b**) LD decay curve illustrates the reduction in LD ($r^2$) with increasing physical distance between markers for each chromosome. (**c**) Gender-based LD decay, where F and M represent female and male samples, respectively. LD decays to half its maximum value at three hundred bp across the genome (refer to Table S7 and S8 for detailed).

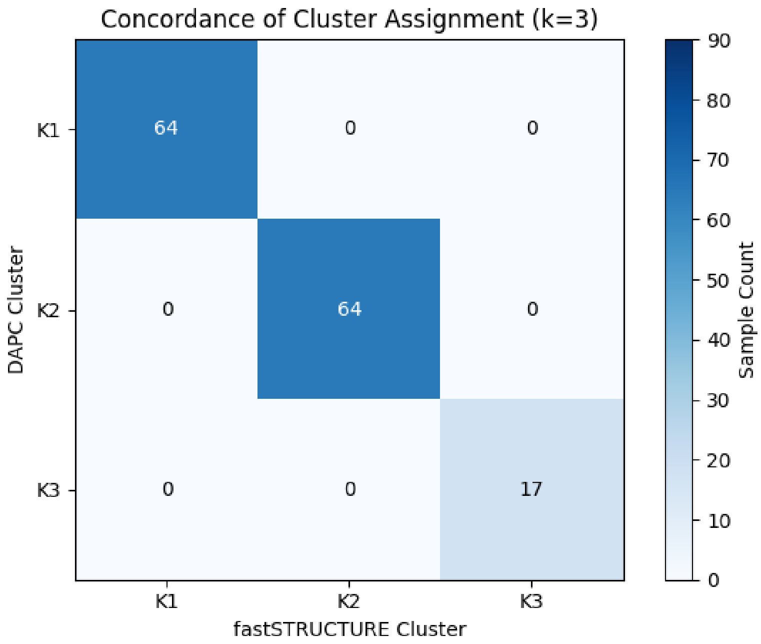


**Figure S3:** Heatmap of the confusion matrix illustrating the concordance between fastStructure and DAPC cluster assignments for K=3.

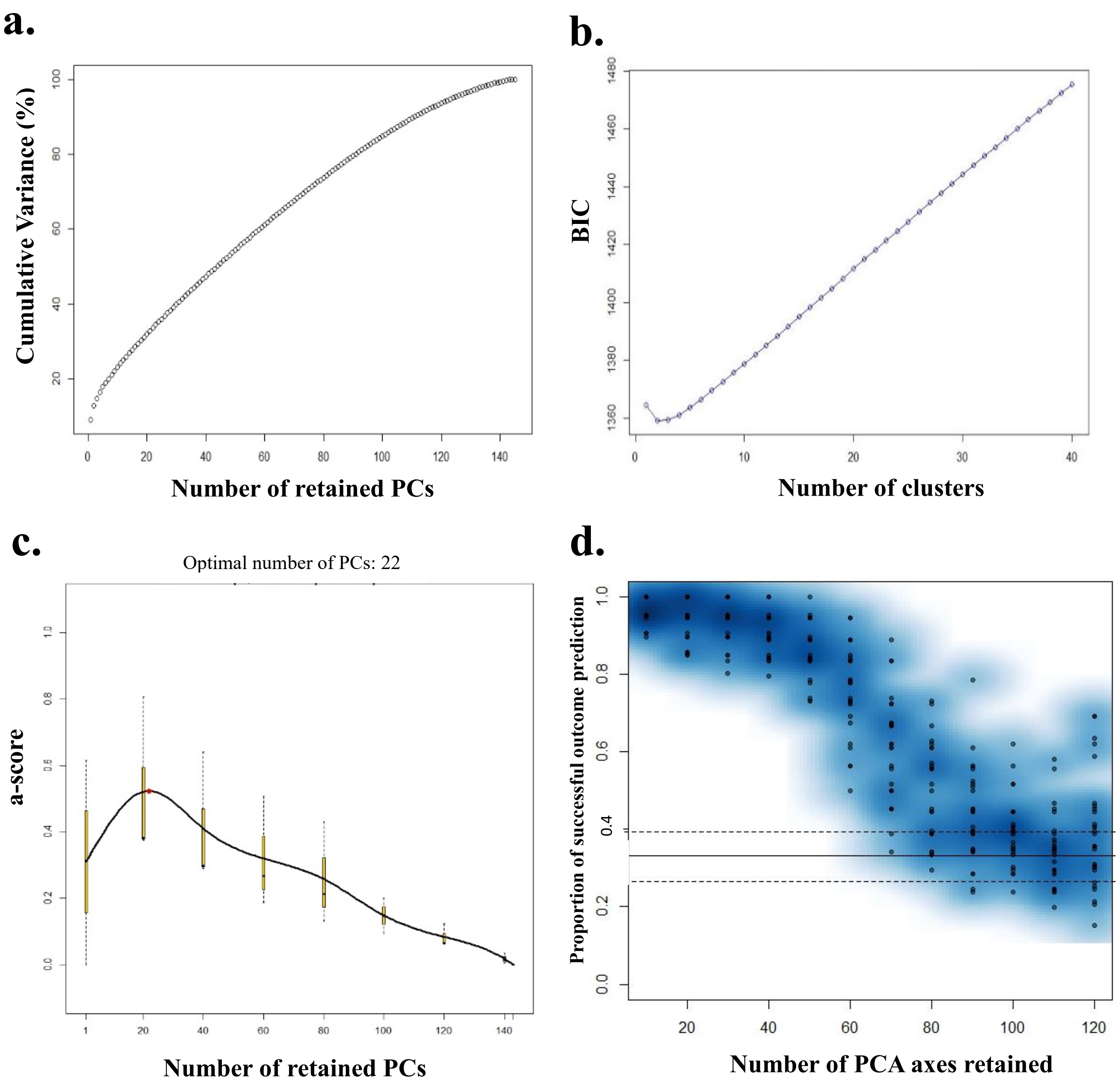


**Figure S4:** Discriminant analysis of principal components (DAPC) performed on 145 *Cannabis* landrace accessions using 233K high-quality SNPs. (**a**) Cumulative variance explained by the eigenvalues of PCA. (**b)** BIC values plotted against increasing *k*-values for determining the optimal number of clusters. (**c**) α-score optimization graph, indicating that 22 principal components were optimal for the analysis. (**d**) Cross-validation results from DAPC confirming the optimal number of retained principal components.

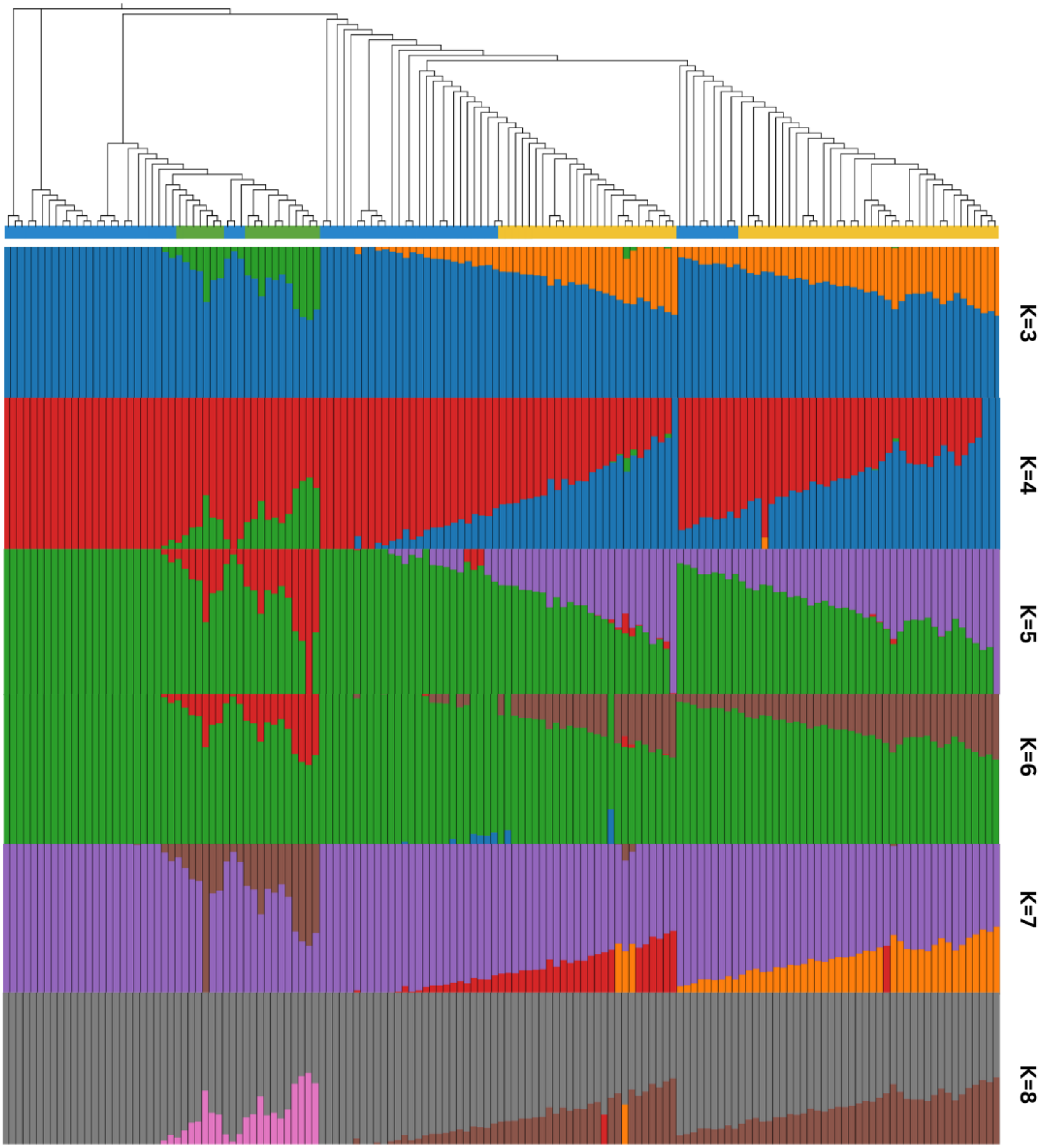


**Figure S5:** Analysis of the population structure of 145 *Cannabis* landrace accessions using the complete set of 233K high-quality SNPs. The admixture plot is shown for *k* values ranging from 3 to 8, generated using fastStructure. The vertical bars represent the landraces, and the y-axis indicates the probability of each individual belonging to a specific subgroup.

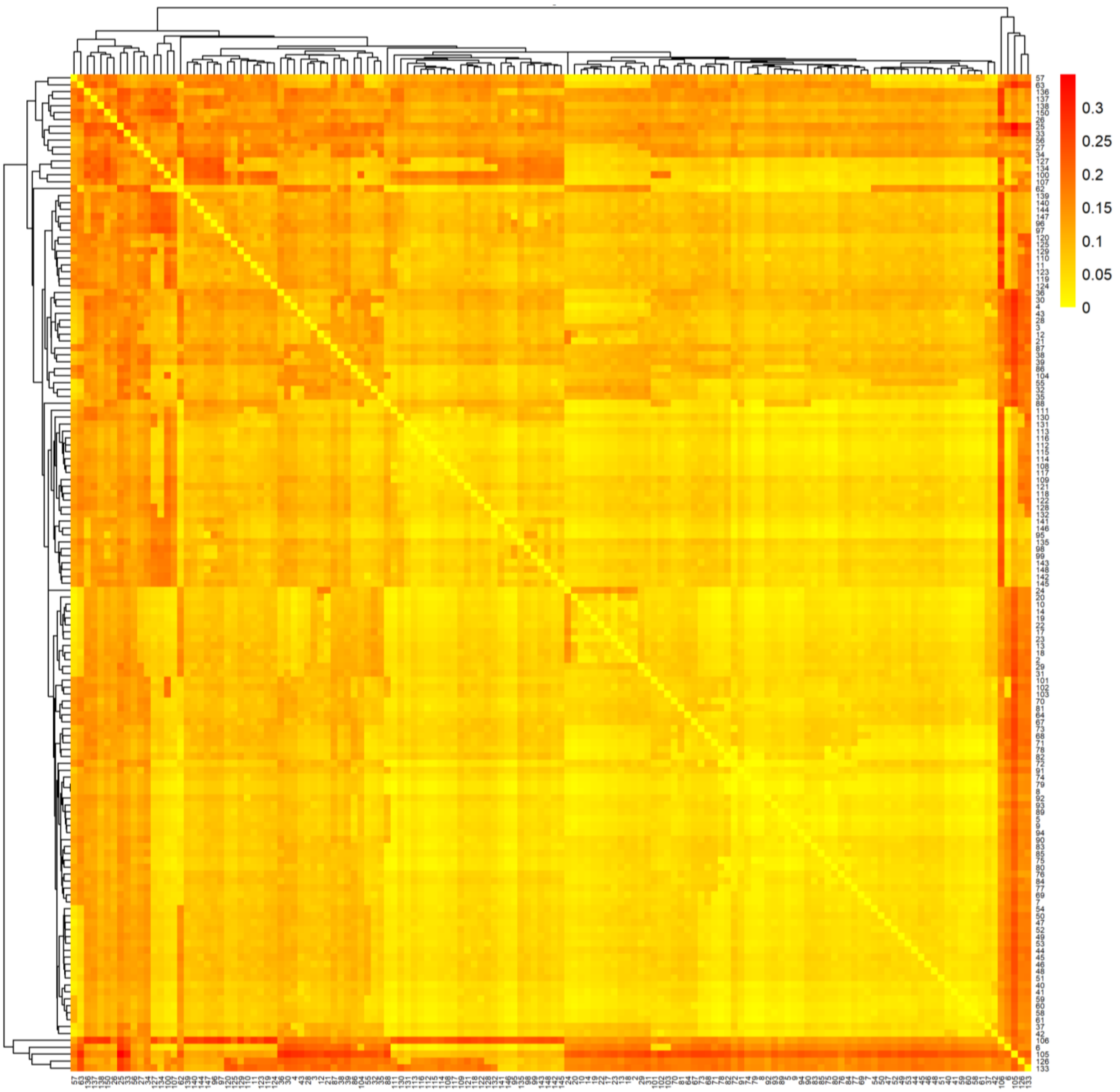


**Figure S6:** Heatmap of pairwise genetic distances among 145 *Cannabis* landrace accessions based on high-quality SNPs. The distances were calculated using the *p-distance* method with gamma-distributed rates among sites and partial deletion of gaps (site coverage cutoff = 95%). Color intensity corresponds to genetic divergence, with darker shades indicating greater distance.

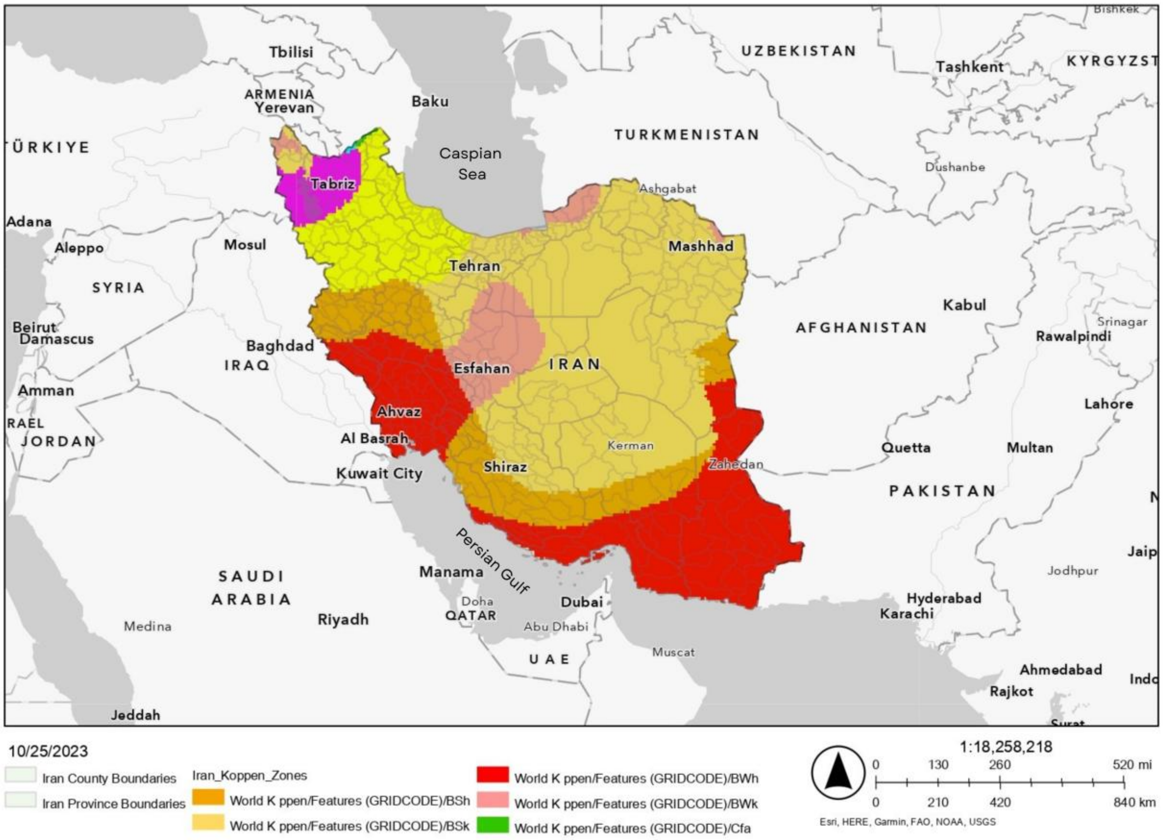


**Figure S7:** A geographical map of Iran marking the five climate zones where the seeds were collected (Adapted from (Babaei et al., 2024)).

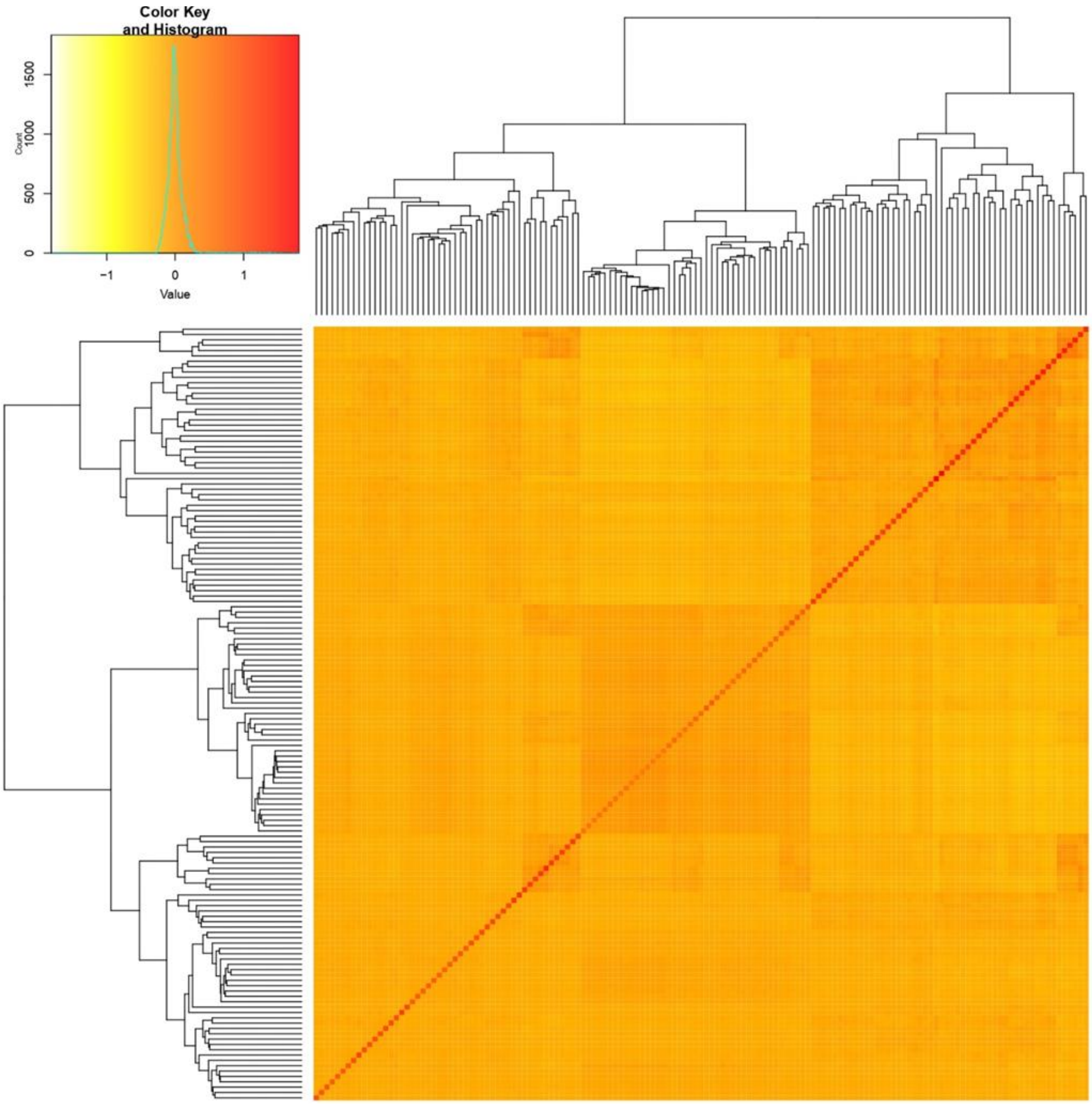


**Figure S8:** Heatmap illustrating the pairwise kinship values among 145 landrace accessions. The color gradient in the histogram reflects the range of co-ancestry coefficients, with darker red tones representing higher genetic relatedness, while lighter yellow shades indicate lower genetic similarity.

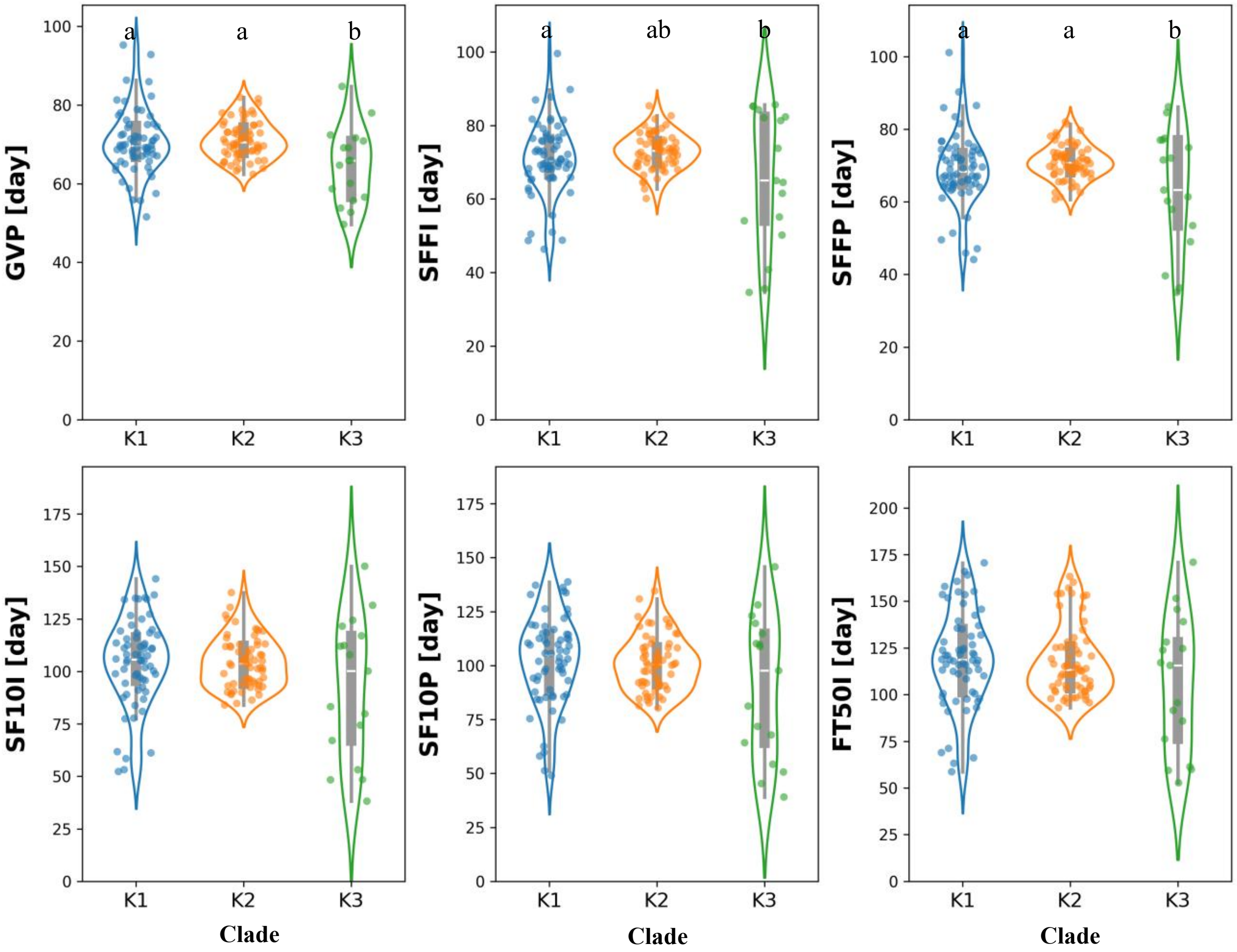


**Figure S9:** Distribution of phenological traits across 145 landrace accessions based on K assignments. The letters show significant differences between clades at the 95% confidence level according to Tukey's HSD test. Abbreviations; GVP: GV Point, SFFI: Start Flower Formation Time in Individuals, SFFP: Start Flower Formation Time in 50% Population, SF10I: Start 10% Flowering Time in Individuals (10% of bracts formed), SF10P: Start 10% Flowering Time in 50% Population (10% of bracts formed), FT50I: Flowering Time 50% in Individuals (50% of bracts formed).

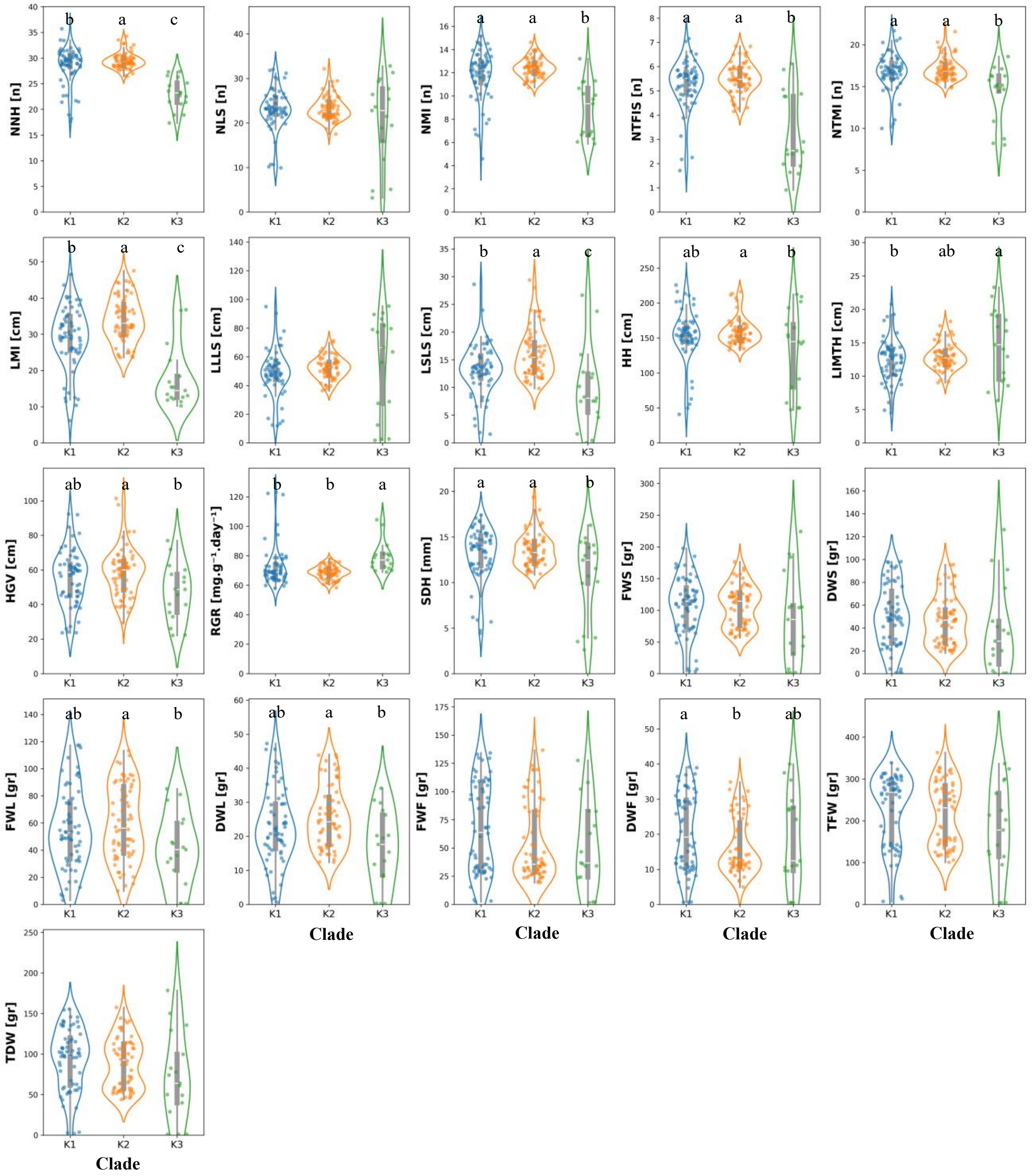


**Figure S10:** Distribution of various morphological traits across 145 landrace accessions based on K assignments. Plots illustrate the phenotypic distribution of traits within clade I (K1), clade II (K2), and clade III (K3). These traits include: Node and Branching Architecture [Number of Nodes on the main stem on Harvest day (NNH), Number of Lateral Shoot (NLS), Number of Nodes on the Main Inflorescence (NMI), Number of Nodes to the First Lateral Shoot (NTFIS), Number of Nodes to the Main Inflorescence (NTMI)]; Growth and Structural Dimension [Length of Main Inflorescence (LMI), Length of Longest Lateral Shoot (LLLS), Length of Shortest Lateral Shoot (LSLS), Height on Harvest day (HH), Length of Internode in the Middle Third of the main stem on Harvest day (LIMTH), Height to GV Point (HGV), Relative Growth Rate (RGR), Stem Diameter on Harvest day (SDH)]; and Biomass Yield [Fresh weight of stems (FWS), Dry weight of stems (DWS), Fresh weight of leaves (FWL), Dry weight of leaves (DWL), Fresh Weight of Flowers (FWF), Dry Weight of Flowers (DWF), Total Fresh Weight (TFW), Total Dry Weight (TDW)]. Different letters above the plots indicate significant differences between clades at the 95% confidence level according to Tukey's HSD test.

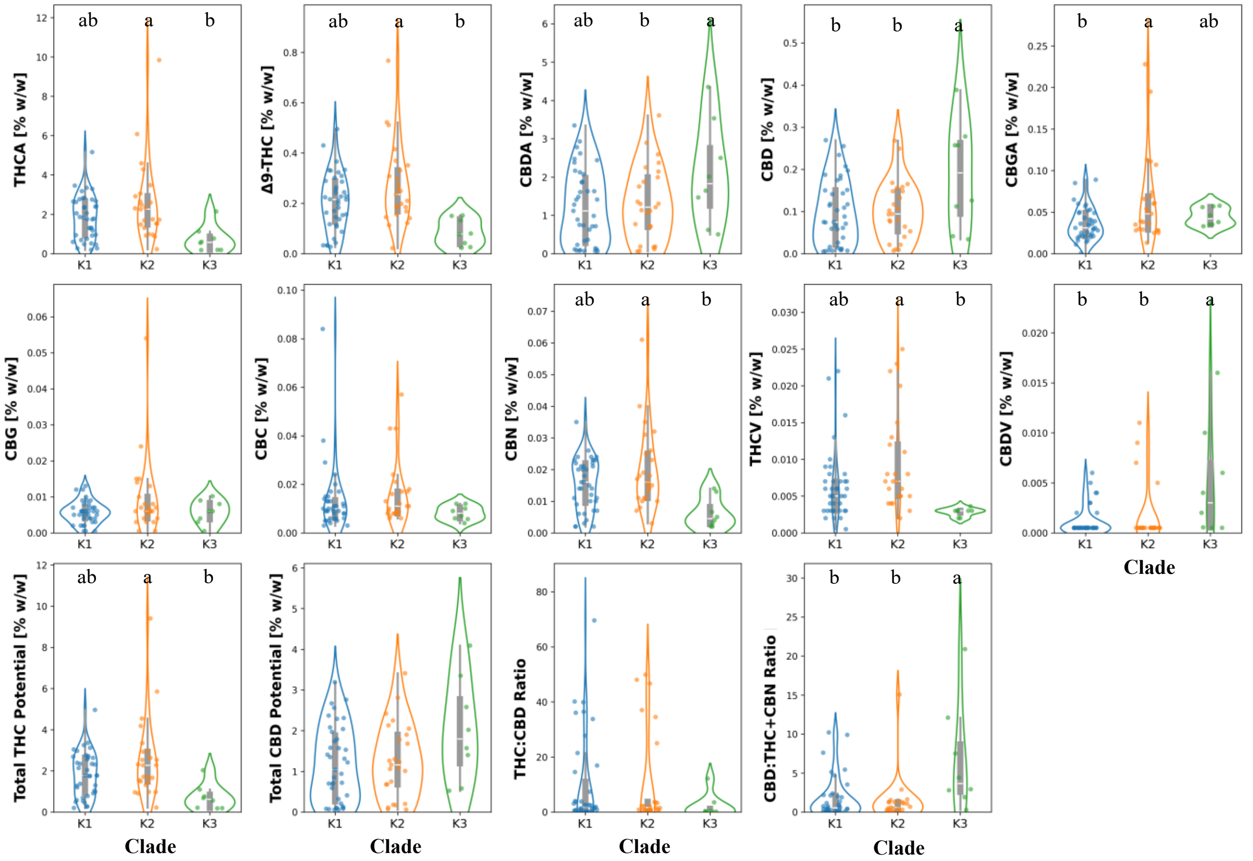


**Figure S11:** Distribution of various phytochemical traits across 145 landrace accessions based on K assignments. Plots illustrate the concentration (% w/w) of Total THC Potential, Total CBD Potential, THCA, $\Delta^9$-THC, CBDA, CBD, CBGA, CBG, CBC, CBN, THCV, CBDV, THC:CBD ratio, and CBD:(THC+CBN) ratio within clade I (K1), clade II (K2), and clade III (K3). Abbreviations: THCA: Tetrahydrocannabinolic acid, $\Delta^9$-THC: $\Delta^9$-Tetrahydrocannabinol, CBN: Cannabinol, THCV: Tetrahydrocannabivarin, Total THC Potential: Potential Tetrahydrocannabinol (Total THC), THC:CBD: Ratio of Tetrahydrocannabinol to Cannabidiol, CBGA: Cannabigerolic acid, CBG: Cannabigerol, CBC: Cannabichromene, CBDA: Cannabidiolic acid, CBD: Cannabidiol, CBDV: Cannabidivarin, Total CBD Potential: Potential Cannabidiol (Total CBD), CBD:THC+CBN: Cannabidiol to (Tetrahydrocannabinol + Cannabinol) Ratio. Different letters above the plots indicate significant differences between clades at the 95% confidence level according to Tukey's HSD test.

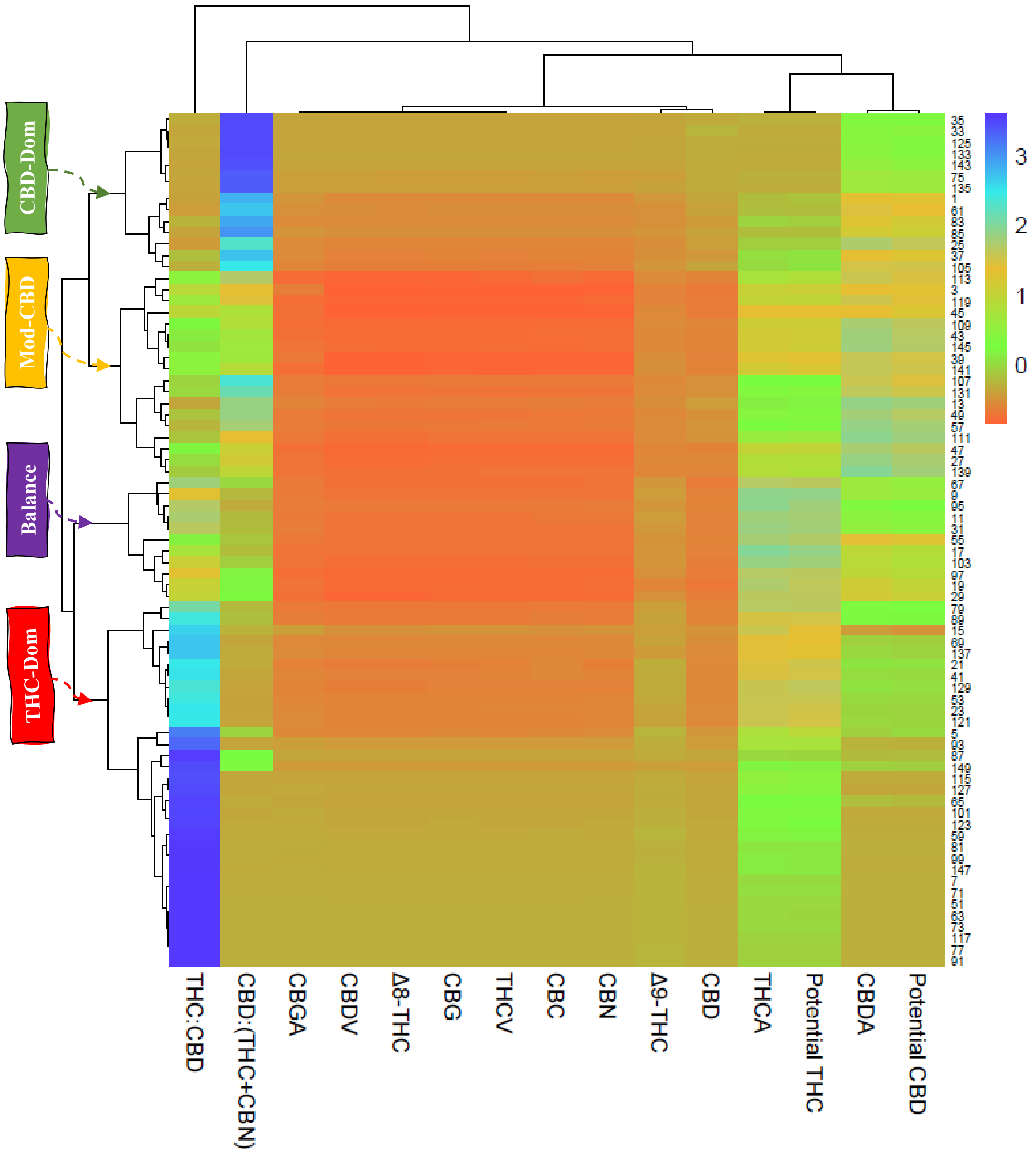


**Figure S12:** Heatmap and dendrogram illustrating the classification and clustering of 145 *Cannabis* landrace accessions based on their phytochemical profiles. The heatmap shows the relative concentrations of various cannabinoids and their derived ratios. The dendrogram on the left indicates hierarchical clustering of accessions based on their phytochemical similarity.

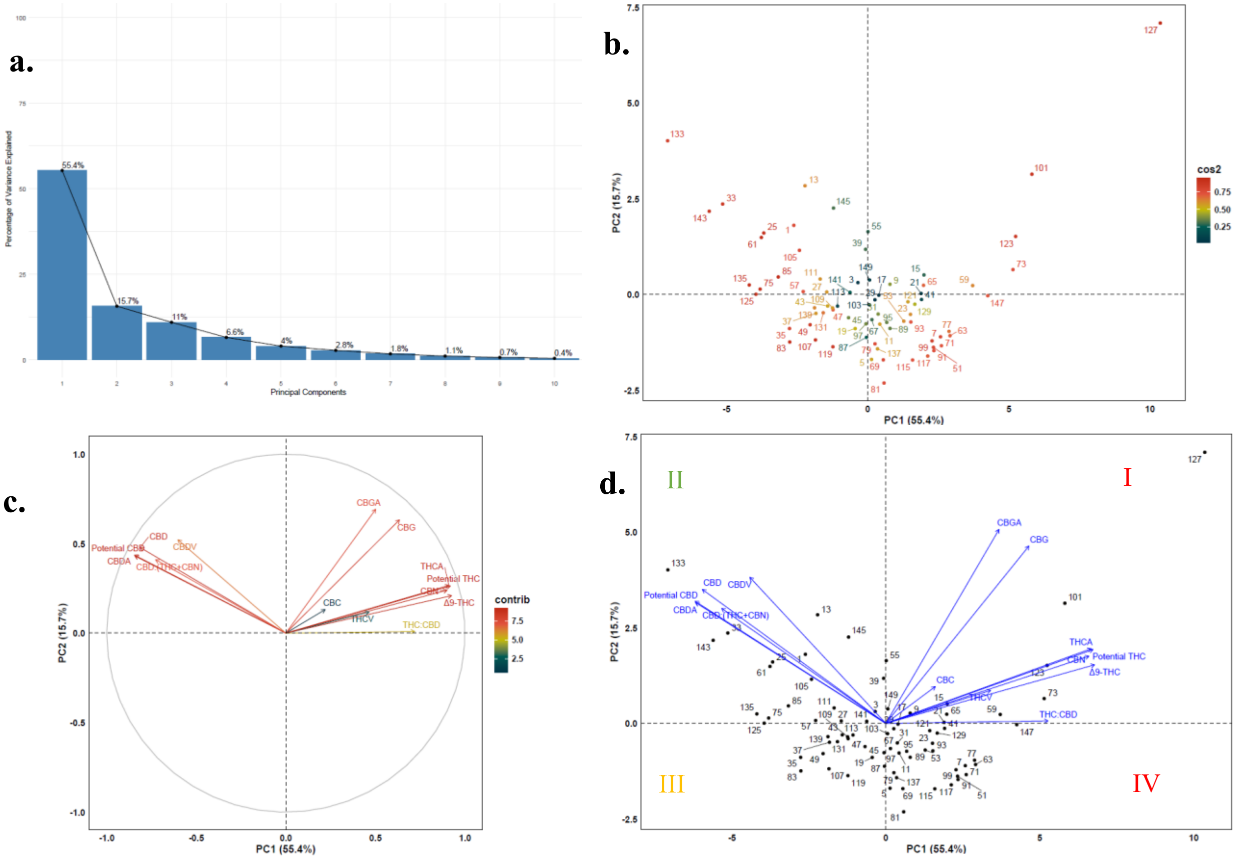


**Figure S13:** Principal Component Analysis (PCA) of phytochemical profiles across 145 *Cannabis* landrace accessions. (**a**) Scree plot showing eigenvalues and the proportion of total phenotypic variance explained by each principal component, illustrating dataset dimensionality. (**b**) PCA scatter plot displaying the distribution of individual accessions in the PC1-PC2 space. Accessions are color-coded by groups, with 95% confidence ellipses illustrating their dispersion. PC1 and PC2 account for 55.4% and 15.7% of the total variation, respectively. (**c**) Biplot of trait loadings, indicating the direction and magnitude of influence of phytochemical traits on PC1 and PC2. (**d**) Phenotypic clustering of accessions based on phytochemical composition. Distinct groupings are associated with key traits: Quadrants I and IV represent THC-dominant (THC DOM) accessions, primarily influenced by CBGA, CBG, THCA, Total THC Potential, CBN, $\Delta^9$-THC, THC:CBD ratio, CBC, and THCV; Quadrants II and III, corresponding to CBD-dominant (CBD-DOM) and moderately CBD (MOD-CBD) accessions, respectively, are both characterized by high levels of CBDV, CBD:(THC+CBN) ratio, CBD, CBDA, and total CBD potential. Additionally, accessions primarily located along the Y-axis of the plot generally represent balanced cannabinoid profiles.

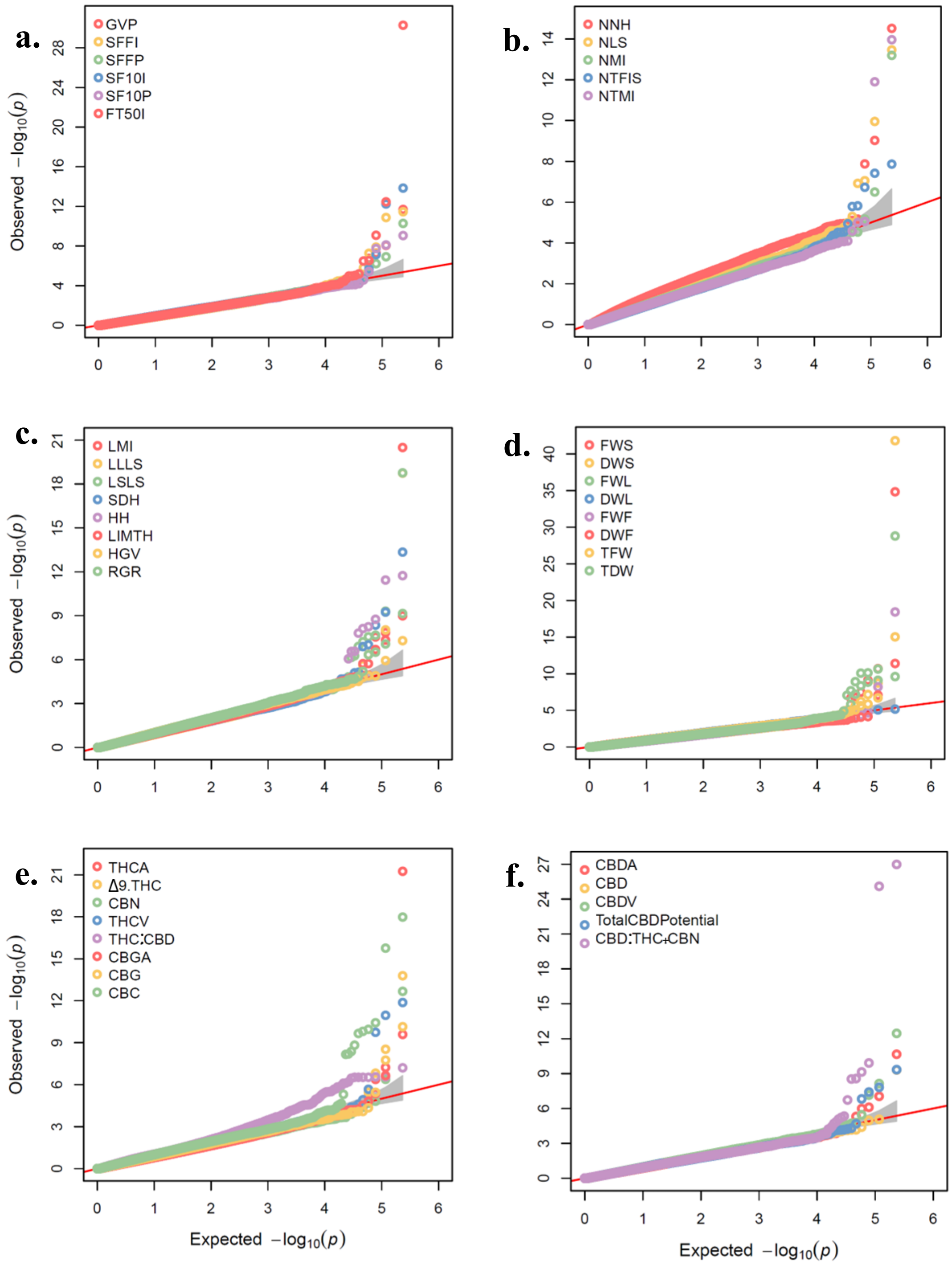


**Figure S14:** Quantile-Quantile (QQ) plots for genome-wide association study (GWAS) results in the 145 cannabis landrace accessions panel. Plots illustrate the distribution of observed versus expected negative logarithm (base 10) *P*-values for: (**a**) Phenological traits, (**b**) Node and Branching Architecture traits (morpho-agronomic), (**c**) Growth and Structural Dimension traits (morpho-agronomic), (**d**) Biomass Yield traits (morpho-agronomic), (**e**) THC-related traits (Cannabinoids), and (**f**) CBD-related traits (Cannabinoids). GWAS was performed using the BLINK method.

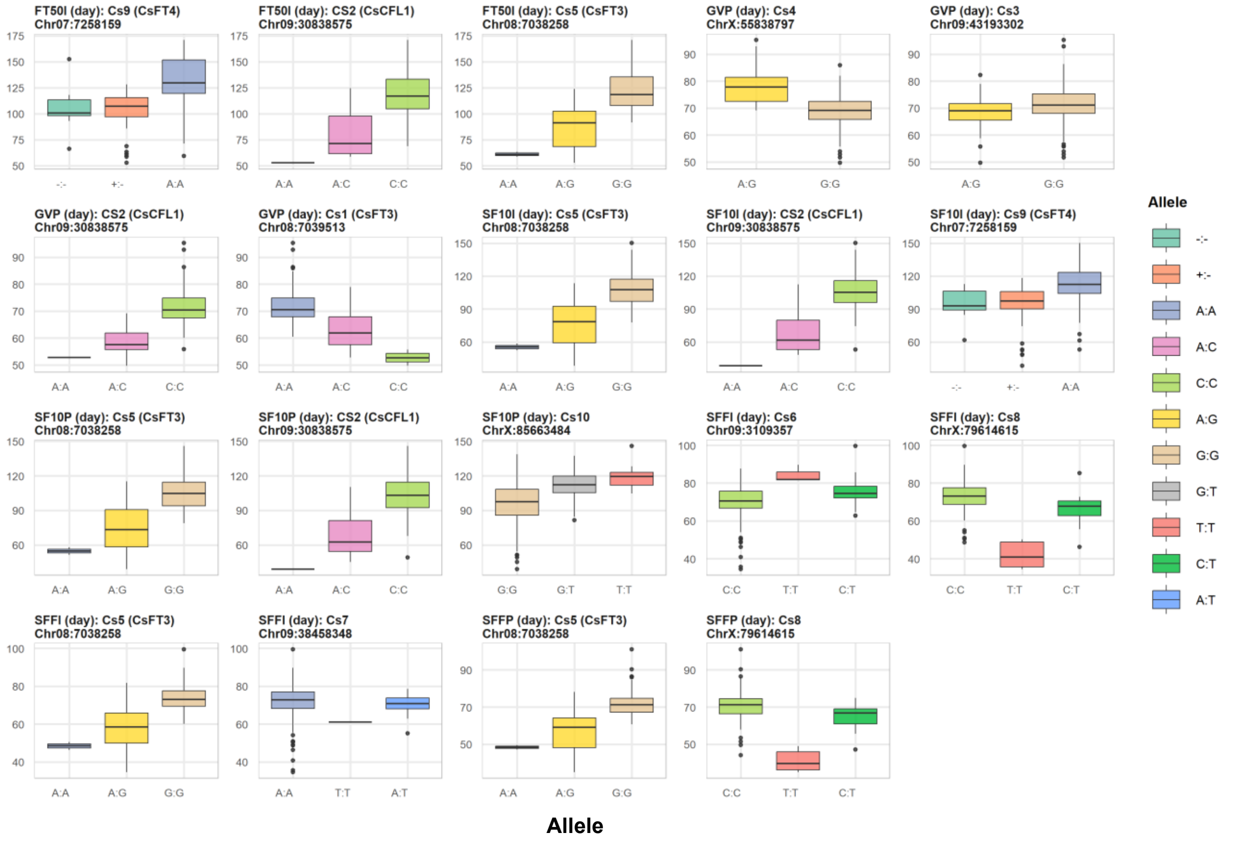


**Figure S15:** Boxplots illustrating the phenotypic effects of different alleles for 10 significant single nucleotide polymorphism (SNP) markers associated with six phenological stages in the 145 cannabis landrace accessions panel. Each panel displays the distribution of phenotype values across various allelic classes for a specific marker. Associations were identified via genome-wide association study (GWAS). Further details are provided in Tables 1 and S14.

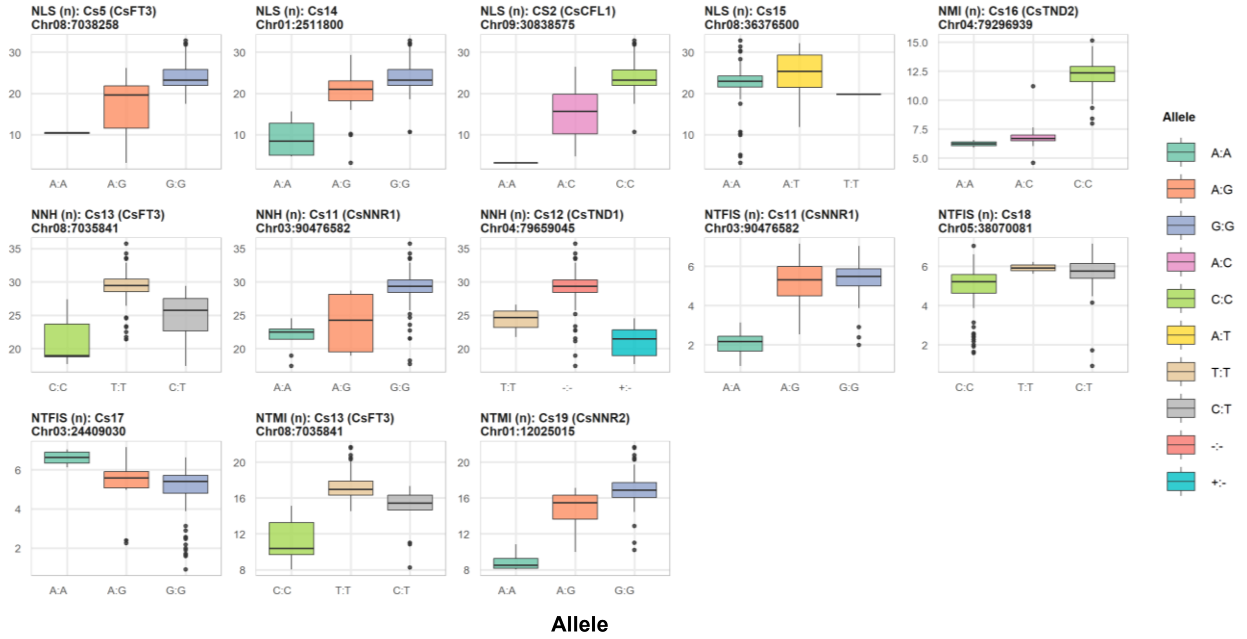


**Figure S16:** Boxplots illustrating the phenotypic effects of different alleles for 11 significant single nucleotide polymorphism (SNP) markers associated with five Node and Branching Architecture (morpho-agronomic) traits in the 145 cannabis landrace accessions panel. Each panel displays the distribution of phenotype values across various allelic classes for a specific marker. Associations were identified via genome-wide association study (GWAS). Further details are provided in Tables 2 and S14.

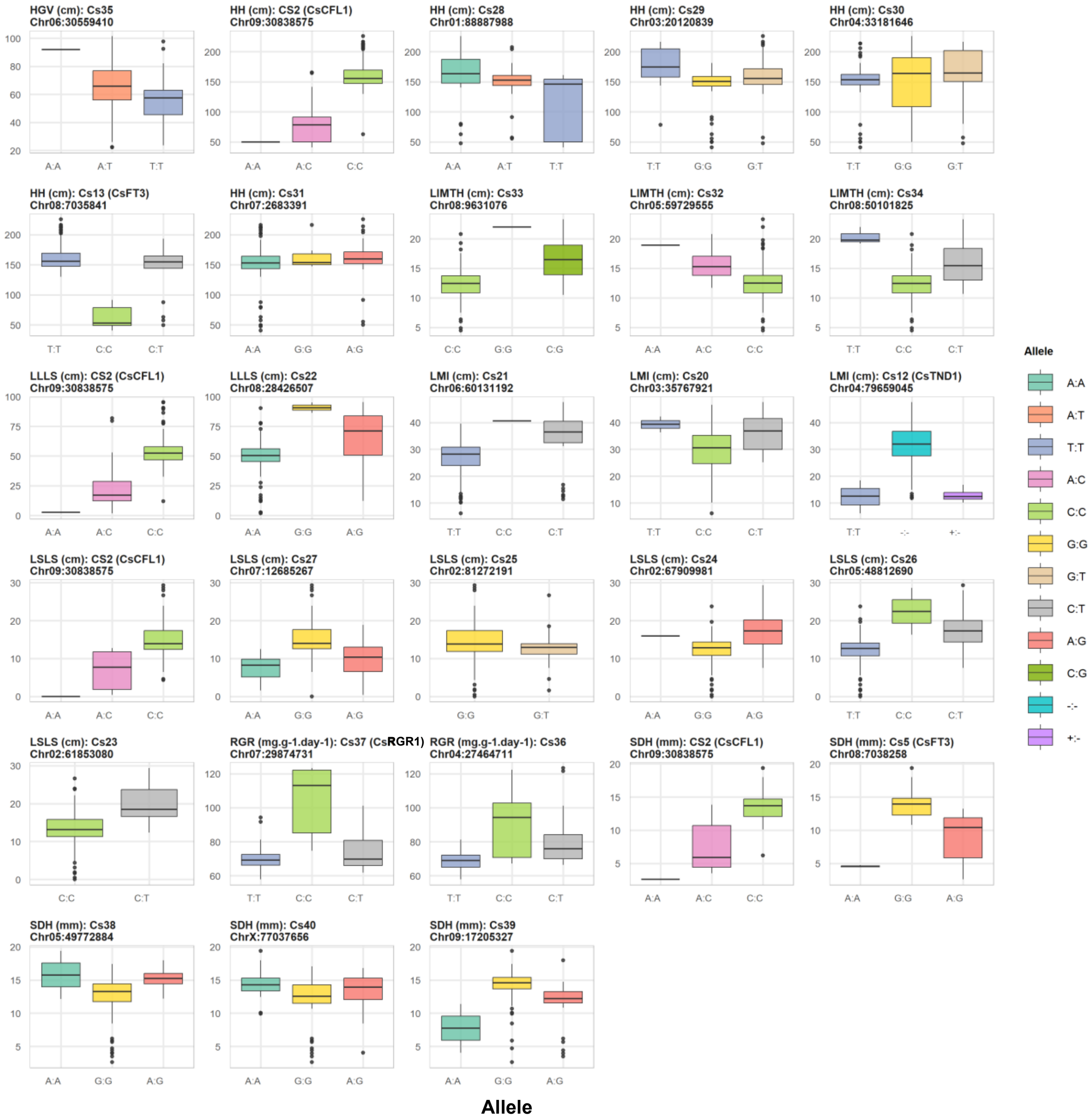


**Figure S17:** Boxplots illustrating the phenotypic effects of different alleles for 25 significant single nucleotide polymorphism (SNP) markers associated with eight Growth and Structural Dimension (morpho-agronomic) traits in the 145 cannabis landrace accessions panel. Each panel displays the distribution of phenotype values across various allelic classes for a specific marker. Associations were identified via genome-wide association study (GWAS). Further details are provided in Tables 2 and S14.

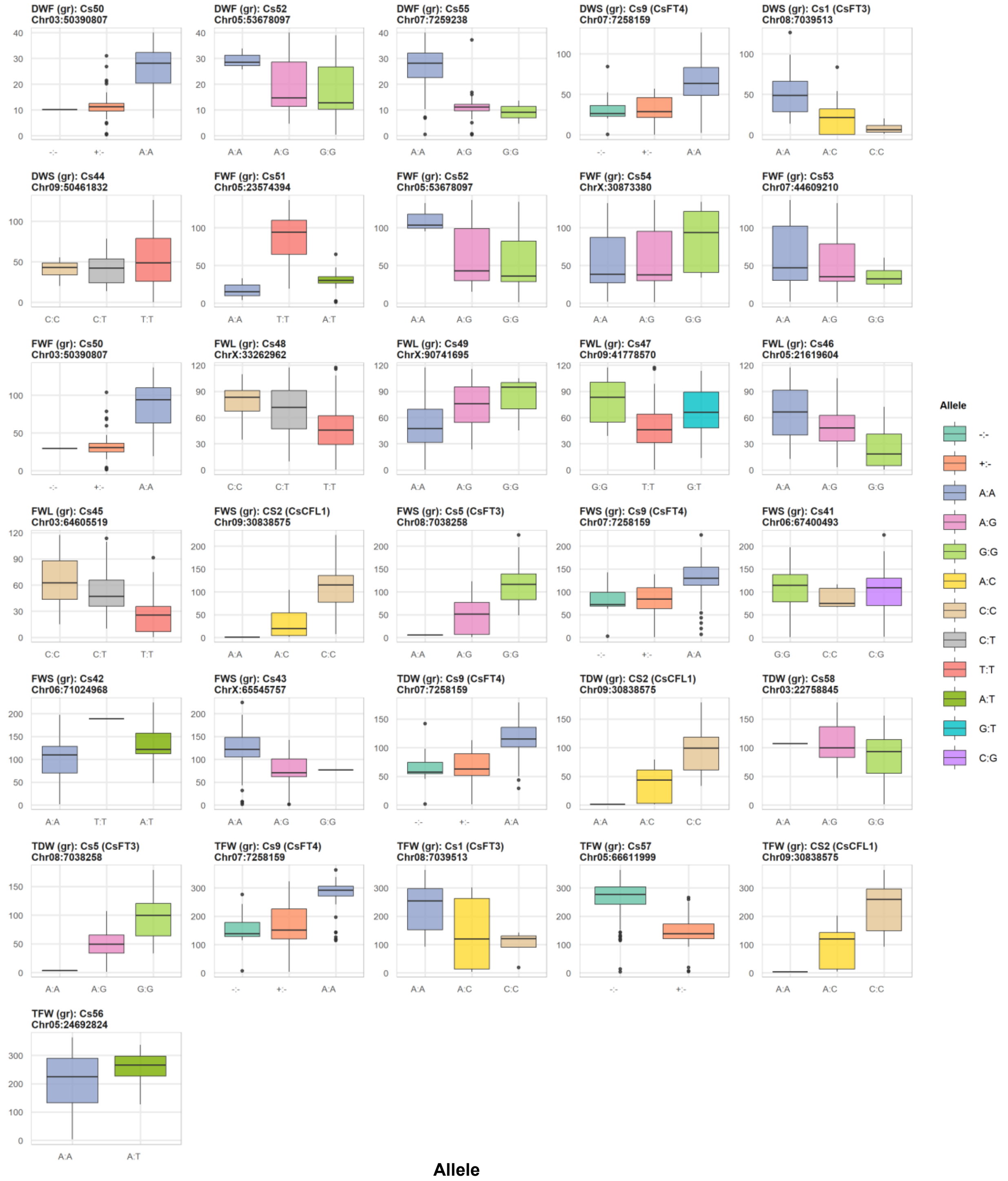


**Figure S18:** Boxplots illustrating the phenotypic effects of different alleles for 22 significant single nucleotide polymorphism (SNP) markers associated with seven Biomass Yield (morpho-agronomic) traits in the 145 cannabis landrace accessions panel. Each panel displays the distribution of phenotype values across various allelic classes for a specific marker. Associations were identified via genome-wide association study (GWAS). Further details are provided in Tables 2 and S14.

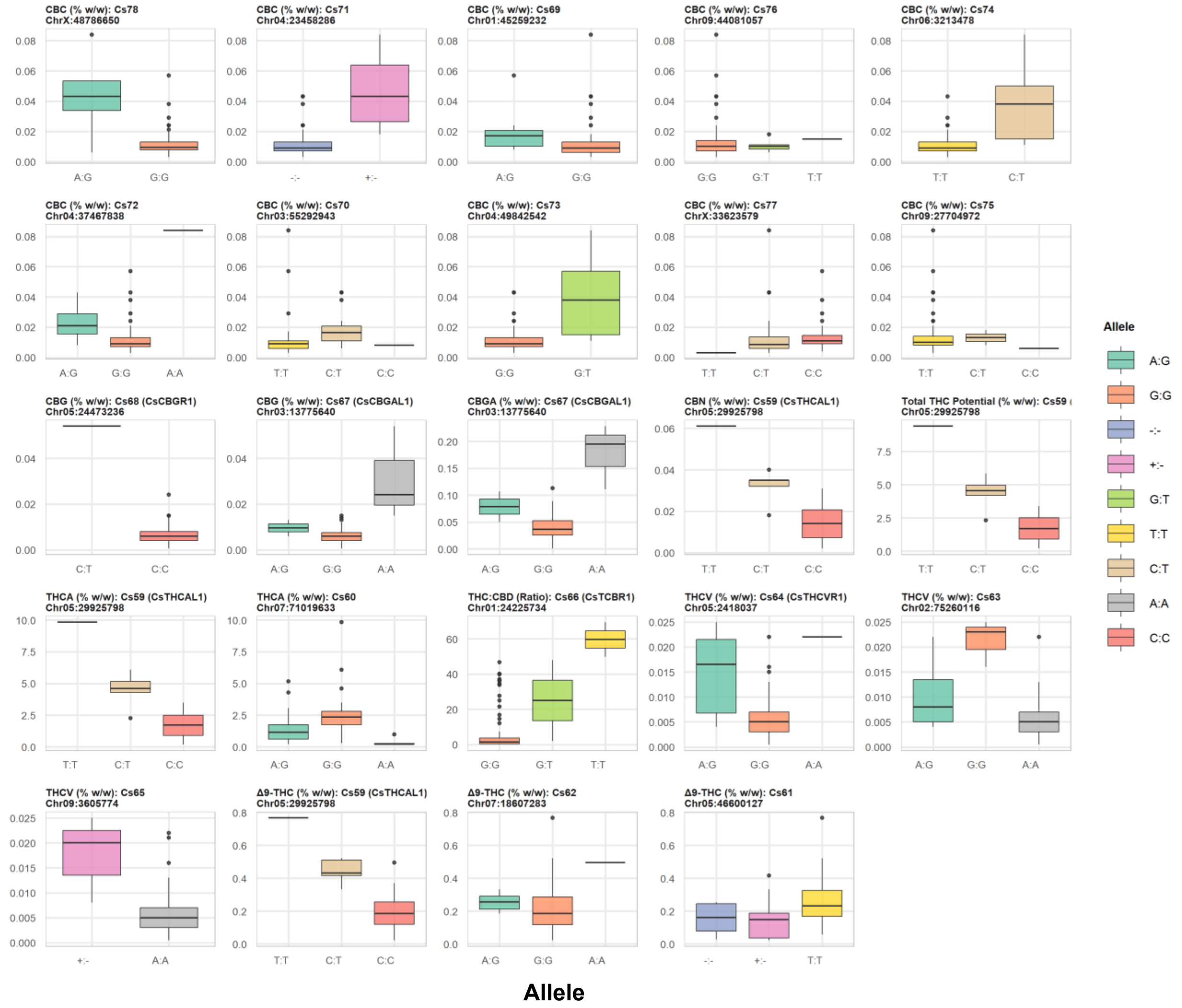


**Figure S19:** Boxplots illustrating the phenotypic effects of different alleles for 20 significant single nucleotide polymorphism (SNP) markers associated with nine THC-related phytochemical traits in the 145 cannabis landrace accessions panel. Each panel displays the distribution of phenotype values across various allelic classes for a specific marker. Associations were identified via genome-wide association study (GWAS). Further details are provided in Tables 3 and S14.

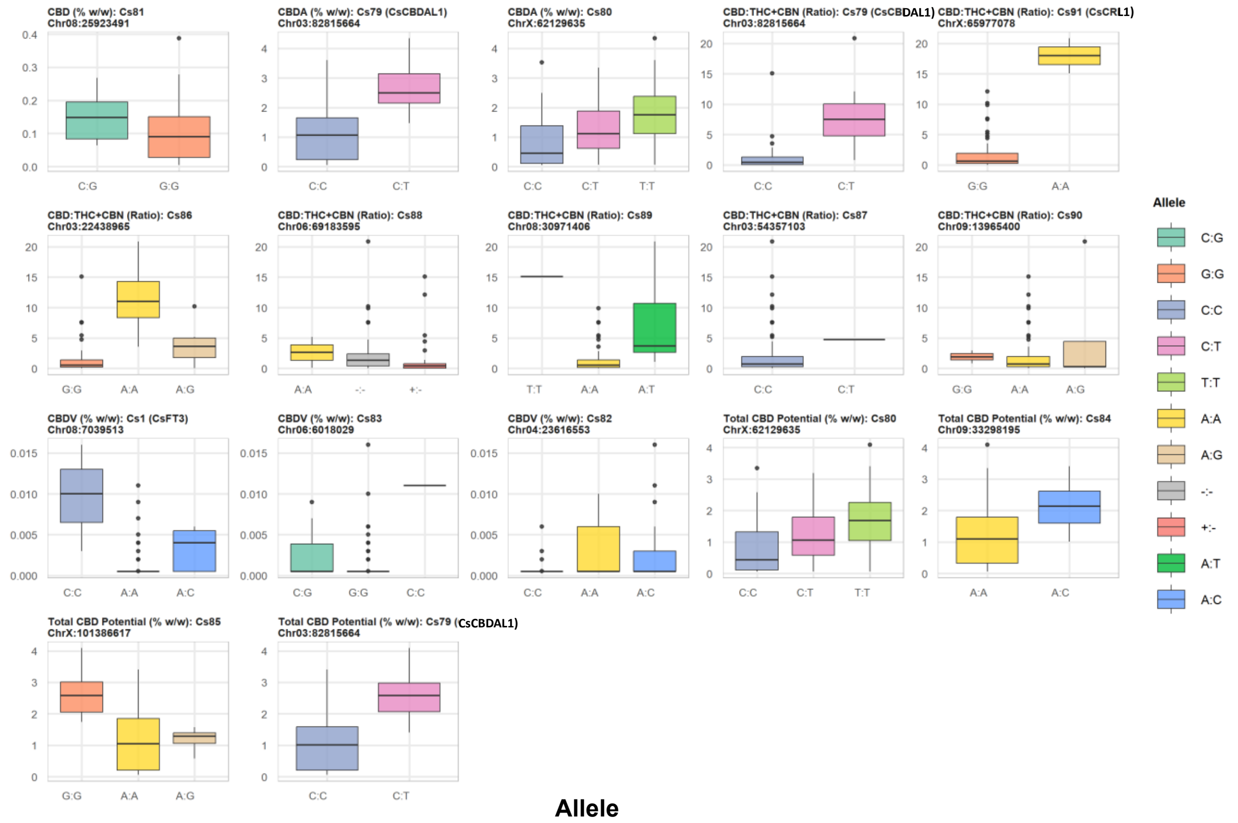


**Figure S20:** Boxplots illustrating the phenotypic effects of different alleles for 14 significant single nucleotide polymorphism (SNP) markers associated with five CBD-related phytochemical traits in the 145 cannabis landrace accessions panel. Each panel displays the distribution of phenotype values across various allelic classes for a specific marker. Associations were identified via genome-wide association study (GWAS). Further details are provided in Tables 3 and S14.
